# Estrogens increase cancer cell efferocytosis to establish an immunosuppressive tumor microenvironment

**DOI:** 10.1101/2024.12.26.630419

**Authors:** Binita Chakraborty, Prabuddha Chakraborty, Michael C. Brown, Daniel Crowder, Aditi Goyal, Rachid Safi, Sandeep Artham, Megan Kirkland, Alessandro Racioppi, Patrick Juras, Debarati Mukherjee, Felicia Lim, Jovita Byemerwa, Kiana Gunn, Scott R. Floyd, Georgia Beasley, Suzanne Wardell, Ching-Yi Chang, Donald P. McDonnell

**Affiliations:** Department of Pharmacology and Cancer Biology, Duke University School of Medicine, Durham, NC, USA; Department of Genetics, University of North Carolina at Chapel Hill, Chapel Hill, NC, USA; Department of Neurosurgery, Duke University School of Medicine, Durham, NC, USA; Department of Radiation Oncology, Duke University School of Medicine, Durham, NC, USA; Department of Surgery, Duke University School of Medicine, Durham NC, USA

**Author notes:** Corresponding author Mailing address: 2138 MSRB3, 3 Genome Court, Box 103057, Durham, NC, USA.

## Abstract

Phagocytic clearance of apoptotic cancer cell_s_ (efferocytosis) by tumor-associated macrophages (TAMs) contributes in a substantial manner to the establishment of an immunosuppressive tumor microenvironment. This puts in context our observation that the female steroid hormone 17β- estradiol (E2) facilitates tumor immune resistance through cancer cell extrinsic Estrogen Receptor (ERα) signaling in TAMs. Notable was the finding that E2 induces the expression of CX3CR1 in TAMs to enable efferocytosis of apoptotic cancer cells which results in the suppression of type I interferon (IFN) signaling. Mechanistically, E2 facilitates calcium-dependent activation of the transcription factor NFATC1, which in turn induces CX3CR1 expression. This drives macrophage polarization towards an immune-suppressive state, increasing the ability of TAMs to engulf pro-inflammatory apoptotic cancer cells. Genetic or pharmacological inhibition of the E2/ER/CX3CR1 axis reversed the efferocytic activity of TAMs, rescued E2-dependent suppression of type I IFN signaling, and potentiated intratumoral adaptive immune cell function. Efferocytosis following radiation-induced cancer cell apoptosis limits the efficacy of radiation therapy. Importantly, we determined that preconditioning with either ER-directed endocrine therapies or CX3CR1 inhibition enhanced the antitumor efficacy of radiation therapy by reversing macrophage suppression and reviving intratumoral T cell activation. Our work defines the mechanisms by which E2 increases the efferocytotic activity of TAMs to establish an immunosuppressive tumor microenvironment and demonstrates how this process can be reversed with endocrine therapies which target ERα.

## Introduction

The primary objective of most of the contemporary immunotherapeutic approaches used to treat patients with cancer is to increase the number and activity of intratumoral T-cells. This can be accomplished using drugs that disrupt the activity of inhibitory receptors on T-cells (e.g., PD1, CTLA4, or LAG3) and/or by delivery of autologous T-cells or engineered CAR-T cells [1, 2]. While these approaches have had a significant impact on outcomes in some cancers, the majority of patients, regardless of tumor type, do not experience durable responses [3]. In many cases, the lack of response to these immunotherapies reflects a low mutational burden within tumors. As important, however, is the ability of cancer cells and/or stromal and immune cells within the tumor to produce factors or establish cell-cell interactions that establish an immunosuppressed tumor microenvironment. Thus, the identification of the factors and cell types that contribute to immunosuppression and development of approaches to mitigate the activities of these processes is a primary focus of drug discovery efforts in this field [4].

Among the most immunosuppressive cells within the tumors are tumor-associated macrophages (TAMs), which can account for up to 50% of the mass of tumors [5]. TAMs can contribute to tumor pathobiology by producing factors that increase (a) angiogenesis, (b) cancer cell proliferation and/or survival and (c) adaptive immune suppression [6–8]. Although the appropriate targeting of TAMs will likely increase tumor immunity, it has proven exceptionally difficult to inhibit the tumor-promoting activities of these immune cells in a specific manner. Systemic depletion of myeloid cells has been explored as an approach with little success likely due to the inhibition of processes that require macrophages to maintain normal homeostasis [5]. There is therefore an unmet need for new therapeutic approaches that can be used to inhibit the immunosuppressive functions of TAMs while sparing the normal function of macrophages. Key to success in these efforts is an understanding the pathways and processes responsible for immunosuppression and the exploitation of this information for new drug development.

A fundamental role of phagocytic myeloid cells (e.g., macrophages and neutrophils) is to clear apoptotic cells throughout the body to ensure tissue homeostasis, moderate inflammatory responses and facilitate tissue repair [9–11]. Myeloid cells accomplish this activity by employing efferocytosis, a process by which they engulf apoptotic cells. If left uncleared, these apoptotic cells would proceed to secondary necrosis, an activity that induces potent inflammatory responses [10]. Phagocytosis and efferocytosis can be distinguished by the fact that the former increases inflammation and stimulates immune responses whereas the latter is primarily anti- inflammatory and immunologically quiescent [10]. Within tumors, macrophages actively engulf and clear apoptotic tumor cell debris through efferocytosis, preventing inflammation that would otherwise activate T cell responses within the tumor microenvironment [12, 13]. Not surprisingly therefore the efficacy of therapeutic interventions that produce apoptotic cells as a byproduct (i.e. chemo/radiotherapeutic regimens) is often limited by increased efferocytosis within the tumor [14–17]. In particular, phosphatidylserine (PtD-Ser) exposed on the surface of apoptotic cells induces debris-stimulated tumor growth and its neutralization has been shown to restore chemo/radiotherapeutic efficacy in breast, NSCLC and melanoma tumors [18–21]. As yet, there are no clinical approaches that can be used to directly inhibit efferocytosis by macrophages. However, some progress has been made in developing antagonists of the fractalkine receptor, CX3CR1, which plays a role in efferocytosis by serving as a “find me” receptor to promote macrophage chemotaxis in response to apoptotic cell-derived fractalkine (CX3CL1) [22].

Previously, we demonstrated that the steroid hormone 17β-estradiol (E2) acting in a cancer cell- extrinsic manner promotes macrophage polarization towards an immune-suppressive state. This in turn leads to CD8^+^ T cell suppression which limits the efficacy of immunotherapies [23]. However, the mechanisms by which these actions of E2 are accomplished have not been established. Thus, in this study we probed the mechanism(s) by which E2 impacts macrophage function, revealing a pathway by which this hormone increases intracellular calcium, resulting in the activation of the transcription factor NFATC1, which in turn increases the expression of CX3CR1. Expression of CX3CR1 increased the engulfment of apoptotic cancer cells, suppressed type I IFN signaling leading ultimately to the suppression of T cell activity. Importantly, genetic or pharmacological inhibition of either the ER or CX3CR1 regulated signaling pathways attenuated efferocytosis, reversed the suppression of myeloid cell intrinsic interferon signaling, and promoted CD8^+^ T cell responses in tumors. The potential near-term clinical impact of this work is significant given the availability of several endocrine therapies that are approved for clinical use.

## Results

### E2 suppresses myeloid cell-intrinsic type I interferon signaling

Recently we demonstrated in mouse models of several different solid tumors that E2-activated ERα signaling in macrophages suppresses CD8^+^ T cell activation resulting in increased resistance to immune checkpoint inhibitors (ICIs) [23]. However, the mechanisms by which E2/ER accomplishes these activities, as well as how and if they could be manipulated therapeutically remain to be determined. To address this question, we used the Lewis Lung Carcinoma (LLC1) as a model system. Tumor cells were subcutaneously injected into ovariectomized mice and supplemented with either placebo or E2. Ovariectomy was used to eliminate the confounding influence of cyclical gonadal production of estrogens (and progestins). The E2 dosing regimen resulted in the expected increase in uterine weights in animals compared to their ovariectomized counterparts (**Supplemental Fig 1A**), confirming adequate exposure to this hormone. E2 promoted the growth of LLC1 tumors, similar to what we observed in murine models of melanoma [23]. Importantly, tumor growth was completely blocked by genetic deletion of ERα in myeloid cells (*Esr1*^f/f^; LysMCre) confirming that the effects of estrogens in this model were occurring in a cancer cell extrinsic manner (**Figure 1A**). These data validate the utility of this murine model to address the mechanisms by which E2/ER- signaling in myeloid cells impacts tumor pathobiology. To determine how the E2/ERα axis influences myeloid cell gene expression within tumors, we isolated intratumoral CD11b^+^ myeloid cells from either placebo or E2-treated LLC tumors and performed mRNA sequencing. Differential expression analysis yielded 708 downregulated genes (>2 fold; p-adjust<0.05) and 175 up-regulated genes (>2 fold; p-adjust<0.05) upon E2 treatment (**Figure 1B**). Gene ontology (GO) analysis indicated that E2 treatment resulted in a significant downregulation of pathways associated with type I interferon (IFN) signaling (**Figure 1C-D**) but increased the transcripts associated with calcium ion transport (**Figure1C**). The expression of candidate genes was confirmed using qRT-PCR in an independent study performed using myeloid cells isolated from LLC tumors treated with or without E2 (**Supplemental Figure 1B**). Similarly, E2 treatment suppressed IFN induced gene expression in the A7C11 mammary tumor model, where we have also demonstrated that cancer cell-extrinsic actions of E2 result in increased tumor growth [24] (**Supplemental Figure1C)**.

**Figure 1.**
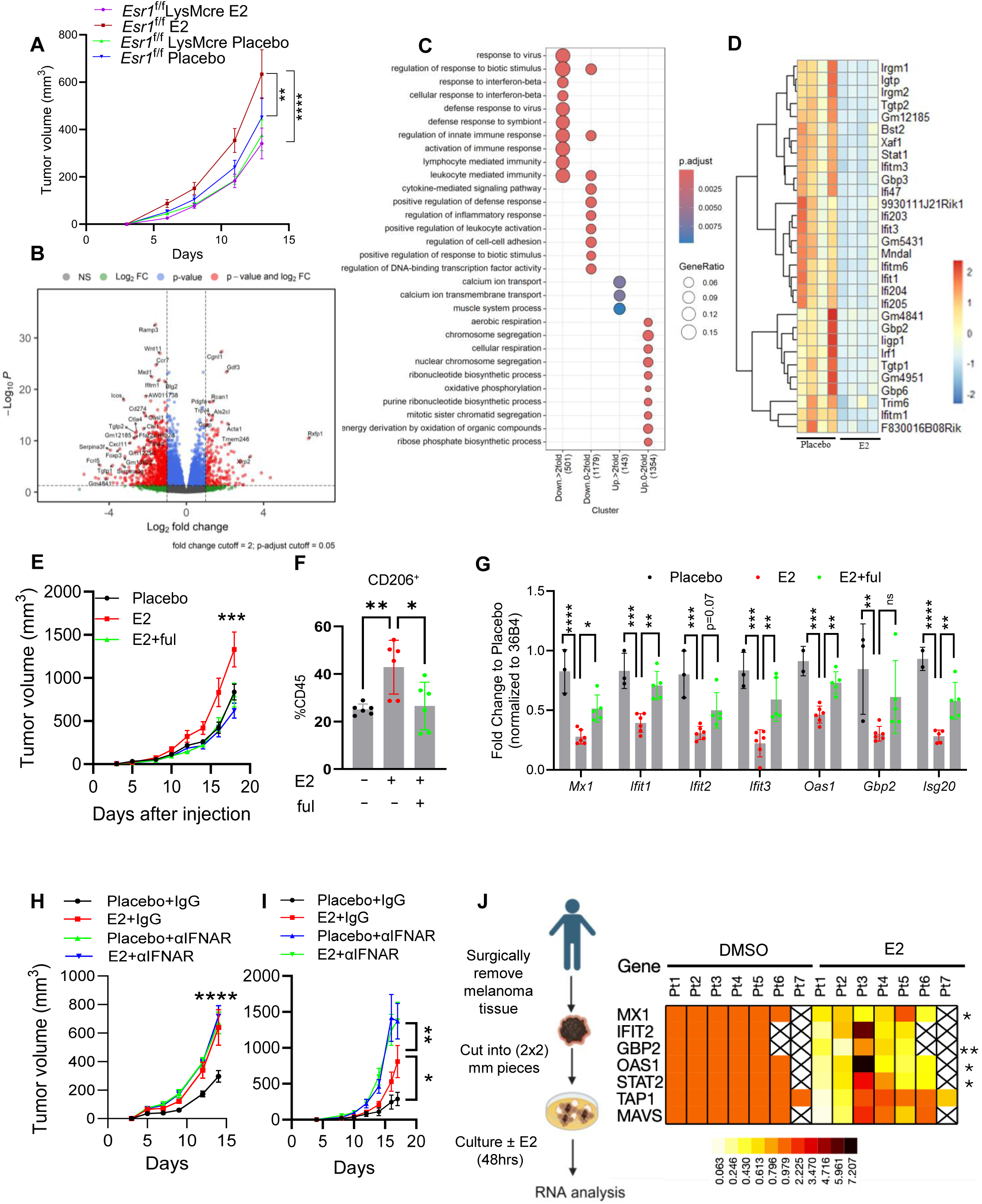
E2 suppresses myeloid cell-intrinsic type I interferon signaling. Syngeneic tumor growth of LLC1 (0.5 x10^5^) cells injected in ovariectomized *Esr1*^f/f^(n=10) and *Esr*1^f/f^; LysMCre (n=10) mice treated ± E2 (**A**). Volcano plots demonstrate genes that are differentially expressed in myeloid cells ± E2 (**B**). GO pathway analysis of RNA-sequencing data performed on intratumoral myeloid cells (Live dead^neg^ CD45^+^, CD11b^+^) isolated from LLC1 tumors injected in ovariectomized mice ± E2 (n=4/group) (**C**). Heatmap showing expression of interferon-stimulated genes (ISGs) as deduced from RNA seq described in figure 1B (**D**) Syngeneic tumor growth and accumulation of intratumoral CD206+ macrophages in LLC1 tumors injected in ovariectomized mice treated ± E2 and E2+fulvestrant (100mg/kg/5 days- intramuscular injection) (**E-F**). Expression of ISGs in intratumoral myeloid cells isolated from LLC1 tumors treated with E2 and E2+fulvestrant (n=5-6 mouse/group) (**G**). Primary tumor growth of LLC1 and B16F10 injected in ovariectomized C57BL6 mice ± E2 in presence or in absence of blocking antibody against α-IFNAR (500µg/mouse/ 3 days-intraperitoneal injection) (n=10 mouse/group) (**H-I**). Scheme and expression of ISGs in melanoma tissues resected from patients and ex vivo treated ± E2 (1nM-48hrs) (**J**). Data represents S.E.M. Significance is calculated by Students t-test, one-way ANOVA, or two-way ANOVA followed by Tuckey’s multiple correction (*p<0.05, **p<0.01, ***p<0.001, ***p<0.0001.) Schematics in J were created by biorender.com.

Gene expression signatures were developed using the human orthologs of the top 50 up- regulated genes (p<0.05) or top 50 down-regulated genes (p<0.05) that we identified in murine macrophages. These were used to query the TCGA database for their association with survival in tumors that were classified as TAM enriched (TAM enrichment score calculated xCell -gene signature-based method) [25]. This analysis revealed that “E2- downregulated genes”, i.e. those that are expressed higher in placebo relative to E2 treated TAMs, were associated with improved overall survival in multiple cancer types (Bladder Cancer, Breast Adenocarcinoma, Lung Adenocarcinoma, Ovarian Cancer, Sarcoma, Uterine Corpus Endometrial Carcinoma). Conversely, E2 upregulated genes, i.e., genes whose expression was lower in the placebo condition, were associated with moderately shorter survival only in ovarian cancer and sarcoma (**Supplemental Figure 2 (A-L**)). Due to (i) the robust association of the E2-downregulated gene signatures with survival, (ii) the observation that the type I IFN signaling pathway was among the most significantly impacted by E2 and (iii) the established role of type I IFN signaling in cancer immune surveillance [26], we next sought to define mechanisms by which estrogens suppress IFN signaling.

Having established a role for estrogens (E2) in regulating myeloid cell intrinsic ISG expression, we further interrogated whether pharmacological inhibition of estrogen signaling alters the suppressive effects of E2 on the expression of ISGs. To address this question, we established LLC tumors in ovariectomized C57BL/6J mice that were treated with either placebo, E2, or E2 in combination with the <u>S</u>elective <u>E</u>strogen <u>R</u>eceptor <u>D</u>ownregulator (SERD), fulvestrant. In this model, treatment with E2 induced the expression of the immunosuppressive marker, CD206^+^, in TAMs, in a manner that was reversed by the addition of fulvestrant (**Figure 1E-F)**. Importantly, it was determined that E2-mediated suppression of ISGs in intratumoral myeloid cells could be reversed by treatment of the mice with fulvestrant (**Fig 1G**).

IFN signaling promotes myeloid cell activation, antigen presentation/antitumor T cell priming, and T cell migration- and activation-processes which we and others have demonstrated to be negatively impacted by E2 signaling in the tumor microenvironment [23, 27–30]. Thus, we probed the functional importance of E2-dependent inhibition of interferon signaling in the tumor growth phenotypes observed. To this end LLC tumors were established in ovariectomized mice treated with or without E2 with concomitant addition of IgG (control) or a blocking antibody targeting the interferon α/β receptors (αIFNAR). Blockade of IFNAR reversed the protective effects of ovariectomy in LLC1 lung tumor models and stimulated tumor growth to the same degree as E2 alone (**Figure 1H**). Similar results were obtained upon blocking the interferon receptors in the B16F10 melanoma model. As in the LLC1 model, IFNAR blockade significantly increased tumor growth in placebo-treated animals. However, it was observed that direct inhibition of the interferon receptors further increased the tumor promoting effect of E2 indicating that in this model some aspects of interferon signaling are not impacted by ER/E2 (**Figure 1I)**.

To confirm the translational significance of our findings in murine models of cancer we evaluated the extent to which E2 suppresses type I IFN signaling in human tumors. Specifically, a fresh tumor tissue fragment assay was performed using freshly resected tumor tissue from patients with melanoma [31]. Isolated tumor fragments were sliced into ∼2mm x ∼2mm x 2mm cubes and were treated *ex-vivo* with either DMSO or E2 for 48 hours. Following treatment, tumor tissues were harvested and a qPCR analysis of the expression of interferon-stimulated genes was performed. Mirroring our findings in murine models (**Figure 1B-C**), IFN-stimulated gene expression (ISG) was suppressed upon E2 treatment in the human melanoma tissue (**Figure1J)**. However, due to the cellular heterogeneity of these tumor fragments, these results may reflect the effect of E2 in decreasing IFN responses from multiple cell types and not necessarily only myeloid cells. Regardless, when taken together, these results highlight the importance of the ER/IFN signaling axis in regulating tumor growth in validated models of solid tumors.

### E2 enhances macrophage efferocytosis

Having established that E2 suppresses IFN signaling in myeloid cells, we next probed the mechanisms responsible for this regulatory activity. For our initial studies, we used RAW 264.7 murine macrophages. In this model, we found that the expression of the ISGs was unchanged by E2-treatment when cells were propagated either in normal media alone, or upon the addition of tumor-conditioned media (TCM) (70% NM + 30% TCM) (Supplemental Figure 3A and 3B). It has been shown by others that the activation of the IFN signaling pathway in TAMs is facilitated by the release of damage-associated molecular patterns (DAMPs). Subsequently, the DNA released by dying cells within the tumor can activate pattern recognition receptor signaling [32–34]. To recapitulate this scenario, we propagated RAW 264.7 cells in the presence of apoptotic splenocytes. Indeed, ISG expression was increased under these conditions which was dramatically reduced by E2 (**Figure 2A**). Similar results were observed when the same assay was performed in the presence of TCM as opposed to NM (**Supplemental Figure 3C**). These data imply that the primary activity of E2 in this system is to limit the chronic inflammation associated with activation of IFN signaling in TAMs that is induced by apoptotic cancer cells.

**Figure 2.**
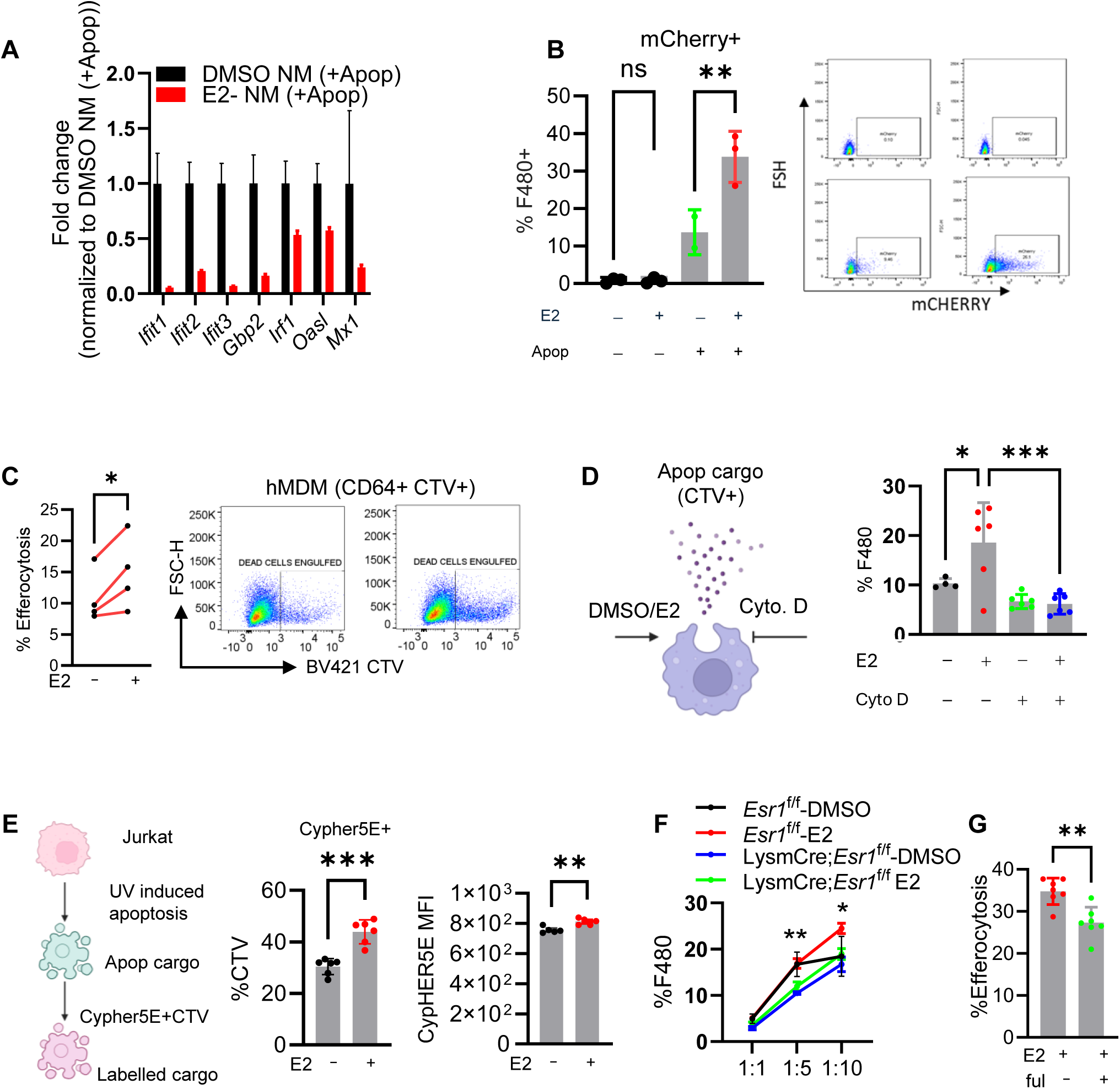
E2 enhances macrophage efferocytosis. Quantitative PCR analysis of interferon-stimulated genes in RAW 264.7 cells that were treated with ± E2 for 72 hrs and were co-incubated with apoptotic cancer cells (LLC) in 1:1 ratio for last 24 hrs (representative of n=2, independent experiments) **(A)**. Flow cytometry quantification of mCherry+J774A macrophages that were treated ± E2 for 72 hrs and co-treated with apoptotic cancer cells (LLC1-mCherry Spectrin) (representative of n=2, independent experiments) **(B)**. Quantification of cell trace violet+ CD64+ human macrophages derived from unidentified donor treated ± E2 (72 hrs) and co-incubated with apoptotic Jurkat cells (1:1) for 1 hr (n=4 independent donor) **(C).** Quantification of Cell-trace violet+J774A macrophages treated ± E2 for 72 hrs. Last 24 hrs the cells were co-incubated with 10nM cytochalasin D followed by co-incubation with cell-trace violet+ apoptotic Jurkat cells for ∼6 hrs (combination of n=2, biological experiment) (**D**). Quantification of number and mean fluorescent intensity of Cypher5E+ BMDM that were treated with ± E2 (48hrs) and co-incubated with CTV+Cypher5E+ apoptotic Jurkat cells (45 mins) (n=3) Quantification of engulfment from BMDM isolated from either *Esr1*^f/f^ (n=3) or *Esr1*^f/f^; LysMCre (n=3) mouse treated with ± E2 for 72 hrs and co-treated with apoptotic (CTV+) Jurkat cells for last 45 minutes at different ratios (1;1; 1:5, 1:10) (**F**). Quantification of percent efferocytosis by J77A macrophages treated ± E2 (72 hrs) and fed with apoptotic Jurkat CTV+ Jurkat cells (**G**). Data represents S.E.M. Significance is calculated by Student’s unpaired t-test (E and G), Student’s paired t-test (C) one-way ANOVA followed by Tuckey’s multiple correction (B and D) and two-way ANOVA followed by Dunnett’s multiple correction (F). *p<0.05, **p<0.01, ***p<0.001, ****p<0.0001). Schematics in D and E were created by biorender.com.

It has been established that TAMs actively phagocytose and dispose of apoptotic cells through efferocytosis, a process which increases immune tolerance in macrophages [12, 35]. Having established that E2 inhibits apoptotic cell-induced expression of ISGs in macrophages, we next explored the possibility that E2 may impact efferocytosis. For this study, we used LLC1 cells engineered to express the fluorescent mCherry-Spectrin fusion protein. Since mCherry is conjugated to Spectrin, it is retained in the apoptotic cells [35], ensuring that any mCherry signal within phagocytes reflects the uptake of cellular material and not released soluble protein (e.g., through micropinocytosis or other processes). Apoptosis in the LLC1-cells was induced using BCL2 and MCL1 inhibitors as previously described [35]. RAW264.7 or J774A cells were then treated with and without E2 in the presence or absence of apoptotic and labeled LLC1 tumor cells for 45 minutes. Macrophages were collected thereafter and the presence of mCherry-Spectrin^+^ cells was quantified by flow cytometry. In this manner it was determined that E2 treatment resulted in a significant increase in the percent of mCherry+ macrophages compared to DMSO treatment, consistent with increased efferocytosis (**Figure 2B and Supplemental Figure 3D**). Similar results were observed when bone-marrow-derived macrophages (BMDM) were co-incubated with cell trace violet (CTV) labeled apoptotic Jurkat cells (**Supplemental Figure 3E**). These studies were extended to human macrophages. Specifically, monocyte derived macrophages (MDMs) were differentiated from PBMCs isolated from four deidentified human donors [31]. Subsequently, the differentiated MDMs were treated for 48 hours with placebo or E2 and then co-incubated with apoptotic CTV labelled Jurkat cells (45 minutes). Evaluation of CTV^+^ macrophages by flow cytometry revealed increased efferocytosis of apoptotic Jurkat cells (**Figure 2C**). To determine whether the changes in efferocytosis observed upon E2 exposure were indeed due to enhanced engulfment capacity, we cotreated macrophages with cytochalasin D- an actin polymerization inhibitor. As expected, treatment with cytochalasin D blunted the basal as well as E2-dependent enhanced engulfment of apoptotic cells by J774A macrophages (**Figure 2D**). Thus, the observed increase in CTV within E2-treated macrophages was indeed due to engulfment of apoptotic cells and not to the binding of cellular debris on the surface of macrophages.

We next measured the effects of E2 on the acidification of cargo following engulfment, as failure to acidify and degrade engulfed apoptotic cargo leads to perturbation of cellular homeostasis and promotes inflammation. This is important as others have shown that TAMs actively dispose of engulfed cargo to prevent inflammation [35]. To explore this further, apoptotic Jurkat cells were co-stained with a pH-insensitive dye CTV and a pH-sensitive dye-Cypher5E, where fluorescence of the latter dye is inversely proportional to pH [36]. E2 treatment increased both the number and the fluorescence intensity of cypher5E^+^ macrophages after co-culture with apoptotic Jurkat cells, indicating phagolysosomal activity and degradation of engulfed cargo (**Figure 2E**). We next explored the specific role(s) of ERα in regulating efferocytosis of apoptotic cells by macrophages. Bone marrow derived macrophages (BMDMs) from *Esr1*^f/f^ or *Esr1*^f/f^;*LysMCre* mice treated with and without E2 were incubated with apoptotic CTV-labelled Jurkat cells (1:1,1:5 and 1:10) for 45 minutes. E2 treatment induced efferocytosis in control BMDMs (*Esr1*^fl/fl^) which was substantially reduced in those in which *Esr1* was deleted (LysMCre; *Esr1*^fl/fl^) at 1:10 ratio (**Figure 2F**). Similarly, pharmacological inhibition of E2/ER signaling using the SERD, fulvestrant reduced the efferocytic capacity of macrophages that were exposed to E2 (**Figure 2G**). Together, the results of these studies performed *in vitro* establish a role for E2 in the enhancement of efferocytic clearance of apoptotic cells by macrophages.

### E2 increases efferocytosis by macrophages in the tumor microenvironment to suppress myeloid cell intrinsic type I interferon signaling

Once we have established the role of E2 in promoting macrophage efferocytosis, we next interrogated whether enhanced efferocytosis limits the activation of myeloid cell intrinsic type I IFN signaling. To confirm our findings *in vivo*, we first assessed the phagocytic capacity of peritoneal macrophages after intraperitoneal injection of apoptotic cancer cells into ovariectomized mice supplemented with and without E2 for 3 weeks. This model measures the basal efferocytic activity of macrophages *in situ* without relying on exogenous adjuvants to activate or recruit phagocytes. Apoptotic LLC cells labeled with mCherry-Spectrin were injected intraperitoneally, and peritoneal exudate cells were collected by lavage 45 minutes after injection. Quantification by flow cytometry revealed an increased number of mCherry^+^ macrophages (mCherry^+^F480^+^) macrophages upon treatment with E2 compared to placebo control (**Figure 3A**). Similar results were also observed in TAMs isolated from LLC- mCherry-Spectrin tumors propagated in ovariectomized mice treated with and without E2 (**Figure 3B**). Collectively, these results establish a role for estrogens in promoting efferocytosis of apoptotic cells within tumors.

**Figure 3.**
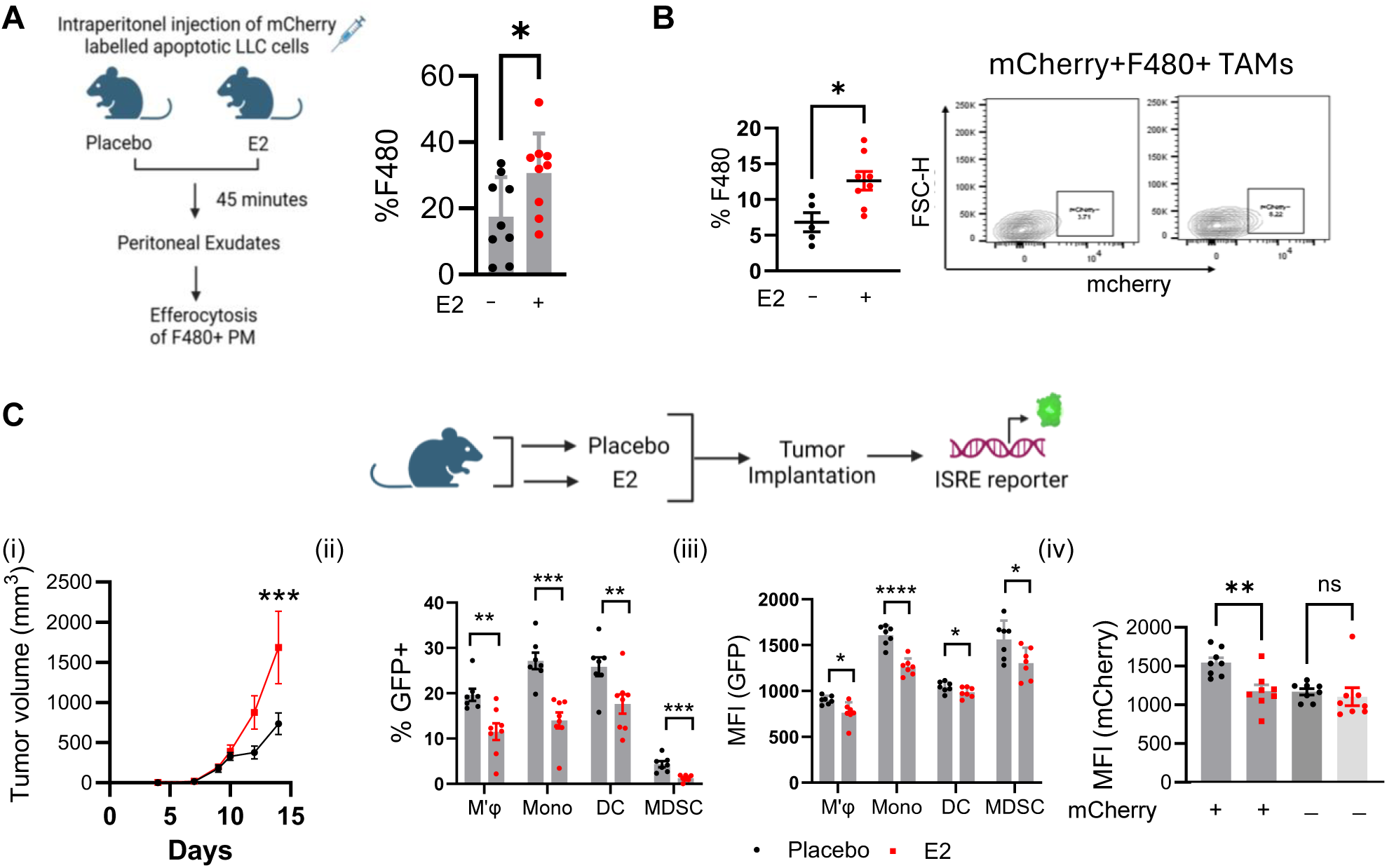
**E2 increases efferocytosis by macrophages in the tumor microenvironment to suppress myeloid cell intrinsic type I interferon signaling**. Quantification of in vivo efferocytosis of peritoneal macrophages from ovariectomized mice treated ± E2 (21 days, n=10 mouse/group) and injected with mCherry-Spectrin+ apoptotic LLC1 cancer cells (4x10^6^cells/mouse) for 45 mins following which peritoneal lavage was collected for flow cytometry analysis (n= 10 mouse/group) **(A)**. Flow cytometry quantification of intratumoral mCherry+ TAMs from LLC1 tumors injected in ovariectomized mice treated ±E2 (n=5-8) and injected with mCherry-Spectrin+ apoptotic cancer cells **(B)**. In vivo tumor growth of LLC1- mCherry-Spectrin cells in ISRE-reporter mouse (Mx1-gfp) (i); quantification of number of GFP+ myeloid subsets (macrophages, monocytes, DC and MDSC) (ii); expression of GFP in different myeloid subsets (macrophage, monocytes, DC and MDSC) (iii); quantification of GFP expression in mCherry+ and mCherry^neg^ intra tumoral myeloid cells (iv) (n=7-8) **(C)**. Data represents S.E.M. Significance is calculated by Student’s unpaired t-test A, B and C (ii) and C (iii), one-way ANOVA followed by Sidak’s multiple correction (C (iv)) and two-way ANOVA followed by Bonferronni’s multiple correction (C (i)). *p<0.05, **p<0.01, ***p<0.001, ****p<0.0001). Schematics in C were created by biorender.com.

Efferocytosis is used by macrophages to decrease the inflammation within cells/tissues and is usurped by TAMs to establish an immunosuppressive environment. Thus, we hypothesized that increased efferocytosis may explain the decrease in IFN signaling observed following treatment with E2. To this end, we used an ISRE reporter mouse (*Mx1^GFP^*) that tracks IFN signaling *in vivo* [37] by expressing GFP under the control of the *Mx1* gene promoter, an established ISG. Ovariectomized Mx1^GFP^ female mice supplemented with either placebo or E2 were engrafted with LLC1-mCherry-Spectrin tumor cells where E2 promoted tumor growth as expected (**Figure 3C (i)**). Analysis of the TAMs from the resultant tumors revealed a significant reduction in the intratumoral accumulation of GFP^+^ myeloid cells (**Figure 3C (ii)**). Further, a significant decrease in the mean fluorescence intensity from GFP in labeled myeloid cells within the tumor microenvironment was observed, indicating a reduction in *Mx1* promoter activity (**Fig 3C** (**iii**)). Importantly, using myeloid cell-intrinsic mCherry fluorescence as a surrogate for LLC1 cell engulfment we identified a specific reduction in GFP fluorescence in mCherry^+^ myeloid cells, but not in mCherry^neg^ myeloid cells (**Figure 3D, (iv)** Taken together, these results Indicate that E2 promotes efferocytosis in TAMs to suppress type I IFN signaling.

### E2 induces the expression of CX3CR1 which results in the suppression of Type I Interferon signaling in myeloid cells

Efferocytosis is a multi-stage process that involves the release of “find me” signals from apoptotic cells thereby facilitating the recruitment of phagocytes [38]. Phagocytes then recognize and respond to “eat me” signals on apoptotic cell membranes, initiating a process that culminates in the phagocytosis and lysosomal elimination of cell debris [39]. Using developmental trajectory analysis of intratumoral myeloid cells isolated from E2-treated tumors, we previously demonstrated an expansion of an ‘M2’-like macrophage cluster with elevated expression of the fractalkine receptor CX3CR1 [23]. CX3CR1 is the cognate receptor for the ‘find me’ signal CX3CL1 that is released by apoptotic cells and promotes efferocytosis by macrophages. Here, we confirmed that E2-treatment led to increased accumulation of CX3CR1^+^ macrophages within the tumor microenvironment *in vivo* (**Supplemental figure 4A**). Moreover, E2 treatment also increased the expression of CX3CR1 on TAMs within the tumor microenvironment and in bone marrow-derived macrophages exposed to TNF (TNFα was used to stimulate basal CX3CR1 expression as described previously [40]) (**Figure 4A and Supplemental figure 4B**).

**Figure 4:**
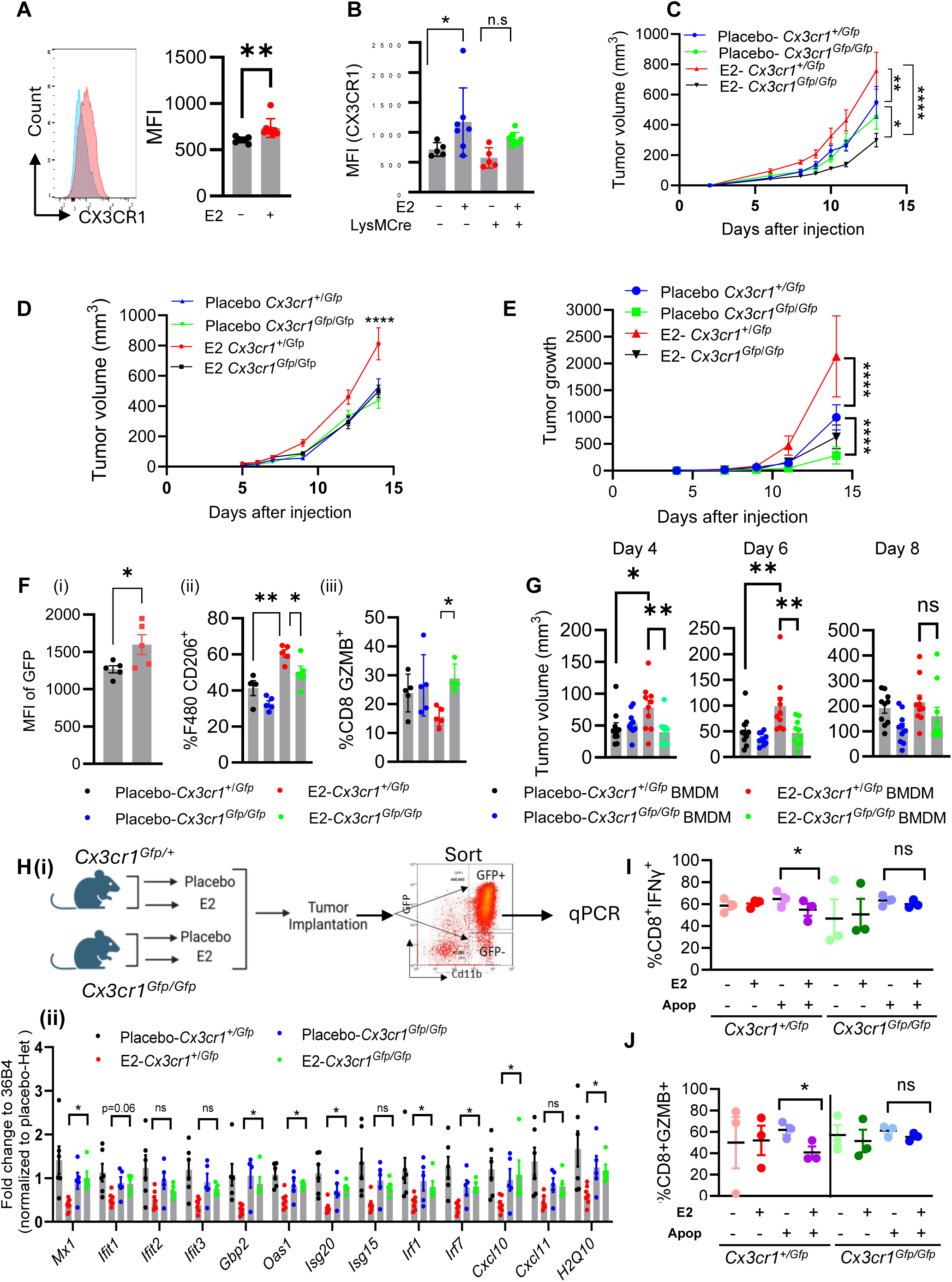
E2 induces the expression of CX3CR1 which results in the suppression of Type I Interferon signaling in myeloid cells. Expression of CX3CR1 in intratumoral TAMs from LLC1 tumors injected in ovariectomized mice treated ± E2 (n=7-8 mice/group) (A). Quantification of mean fluorescent intensity of TAM intrinsic CX3CR1 from ovariectomized *Esr1*^f/f^ and *Esr1*^f/f^; LysMCre mice treated ± E2 and injected with LLC1- mCherry-spectrin cells (n=5-8 mice/group) (B). Syngeneic tumor growth of LLC1 (NSCLC) (n=9-10 mice/group) (C); A7C11 (breast carcinoma) (n=10 mice/group) (D) and B16F10 (melanoma) (n=7-9 mice/group) cells in ovariectomized CX3CR1*^+/gfp^*-HET and CX3CR1*^gfp/gfp^*-KO (null) mice treated ± E2 (E). Quantification of GFP+ TAMs (i); CD206+ TAMs(ii) and GZMB+ CD8+ T (iii) cells from experiment described in figure 3D (F). Quantification of LLC1 tumor volumes when LLC1 tumors were co-mixed with BMDM from CX3CR1*^+/gfp^*-HET and CX3CR1*^gfp/gfp^*-null mice and injected in ovariectomized CD45.1 mouse ± E2 (n=10 mice/group) (G). Schematic representation and quantitative PCR expression of ISGs in sorted CD11b+ GFP+ myeloid cells from LLC1 tumors injected in ovariectomized CX3CR1*^+/gfp^*HET and CX3CR1*^gfp/gfp^*-null mouse ± E2 (n=6-8 mice/group) (H (i) and (ii)). Quantification of IFNγ+(I) and GZMB+ (J) CD8+ T cells when T cells were cocultured with BMDM from CX3CR1*^+/gfp^*-het and CX3CR1*^gfp/gfp^*-null macrophages treated ± E2 (72hrs) in presence or in absence of apoptotic Jurkat cells (last 24 hrs) (n=3 mouse/condition). Data represent mean ± S.E.M. Significance is calculated by unpaired Students t-test (A, F(i)), one-way ANOVA, and pairwise comparison followed by Sidak’s multiple comparisons (I, J and K), Dunnett’s multiple comparisons (F (ii), (iii) and G) (ii)), Bonferronni’s multiple comparisons (B) by or two-way ANOVA followed by Tuckey’s multiple corrections (*p<0.05, **p<0.01, ***p<0.001, ****p<0.0001.)

To determine whether myeloid cell-intrinsic E2/ER signaling drives either the expression of CX3CR1 or the accumulation of efferocytic CX3CR1^+^ TAMs within the tumor microenvironment, we injected ovariectomized mice (*Esr1*^f/f^ and *Esr1*^f/f^; *LysMCre* backgrounds) with LLC1 tumor cells. An increase in myeloid cell-intrinsic CX3CR1 expression in *Esr1*^f/f^ mice was observed in response to E2/ER signaling, which was absent in *Esr1*^f/f^; *LysMCre* mice treated with E2 (**Figure 4B)**. However, ERα depletion did not attenuate the increased intratumoral accumulation of CX3CR1^+^ macrophages upon E2-treatment (**Supplemental Figure 4C**). These results indicate that although recruitment of phagocytic macrophages was mediated by E2 action independent of CX3CR1, myeloid cell-intrinsic ER-signaling was required for the effect of E2 on CX3CR1 expression.

Having established a role of the E2/ER signaling axis in increasing the expression of CX3CR1 on macrophages within the tumor microenvironment, we interrogated the contribution of CX3CR1 to E2-mediated tumor growth. For this purpose, we utilized the *Cx3cr1^+/Gfp^*and *Cx3cr1^Gfp/Gfp^* mouse models. In these mice the EGFP expression sequence replaces the first 390bp of coding exon 2 of *Cx3cr1*, knocking out the endogenous receptor and allowing an evaluation of the expression of CX3CR1 (using GFP expression as a surrogate) in cells in which the receptor is normally expressed [41]. *Cx3cr1*-heterozygous (HET) or *Cx3cr1*-KO (null) mice were ovariectomized, treated with placebo or E2, injected with LLC1 tumor cells, and tumor growth was measured. While E2 treatment promoted tumor growth in the *Cx3cr1* HET mice, homozygous deletion of *Cx3cr1* reduced E2 dependent tumor growth significantly (**Figure 4C**). Similar trends in tumor growth were observed in A7C11- breast carcinoma and B16F10-melanoma models, where E2 increased tumor growth in the *Cx3cr1*-HET background in a manner abolished in *Cx3cr1* null background (**Figure 4D-E**). Collectively, these results indicate that E2 increases the expression of CX3CR1 in TAMs and that this results in an increase in tumor growth.

Analysis of the intratumoral immune-infiltrates revealed that E2 treatment increased CX3CR1 expression (using GFP as a surrogate of *Cx3cr1* expression) in the intratumoral macrophages of *Cx3cr1*-HET mice (**Figure 4F** (**i**)), in a manner that was similar to that observed in WT TAMs (**Fig 3A**). Additionally, homozygous deletion of CX3CR1 led to decreased accumulation of CD206^+^ intratumoral macrophages even in the presence of E2, denoting their shift towards a pro-inflammatory phenotype (**Fig 4F (ii)**). These results indicate that the activity of the recruited macrophages within E2-treated tumor microenvironment is mediated by CX3CR1. This was also associated with reduced intratumoral accumulation of granzyme B (GZMB)^+^ CD8^+^ T cells after E2 treatment in heterozygous mice, which was not observed upon CX3CR1 deletion (**Figure 4F, iii**). However, no changes in overall and exhausted CD8^+^ or CD4^+^ T cells or Tregs were noted upon depletion of CX3CR1 regardless of E2 status (**Supplemental Figure 4D)**.

To further define the role(s) of macrophage-intrinsic CX3CR1 in E2-mediated tumor growth, we performed a co-mixing study. Macrophages that were differentiated from bone marrow progenitors isolated from either *Cx3cr1*-het or *Cx3cr1*-null mice were co-mixed with LLC1 cells and injected into ovariectomized mice treated ± E2. Accelerated LLC1 tumor growth (days 4-6) was observed after LLC: *Cx3cr1*-het macrophage admixture, but not after engraftment with *Cx3cr1*-null macrophages. The initial growth disadvantage of tumors containing *Cx3cr1*-null macrophages was lost after day 8 (**Figure 4G**), likely because of the presence of recipient host- derived CX3CR1^+^ macrophages.

We next explored whether E2-mediated suppression of type I IFN signaling in tumor-associated myeloid cells requires CX3CR1. To this end, GFP^+^ myeloid cells were isolated from LLC1 tumors propagated in ovariectomized *Cx3cr1*-HET or *Cx3cr1*-null mice that were treated with and without E2 (**Figure 4H, (i)**). Deletion of CX3CR1 had a minimal impact on ISG expression in placebo- treated mice. Importantly, qRT-PCR analysis of a large number of ISGs demonstrated that E2- mediated suppression of ISG expression in myeloid cells was rescued upon homozygous deletion of CX3CR1 (**Figure 4H, (ii)**). This suggests that E2 dependent suppression of type I IFN signaling requires CX3CR1. As discussed above, the increased expression of CX3CR1 in macrophages from E2-treated mice was associated with reduced T cell activation (see **Figure 4F (iii)**). Previous studies have demonstrated a causal role for the myeloid cell-intrinsic type I IFN axis in supporting the functionality of T cells in the tumor microenvironment. To explore this further, BMDMs (*Cx3cr1*-HET vs *Cx3cr1*-null) were treated ± E2 ± apoptotic cells and incubated with *in vitro* activated CD8^+^ T cells isolated from spleens of naïve C57BL/6J mice. CD8^+^ T cells incubated with BMDM from *Cx3cr1*-HET mice+E2 (and with apoptotic cells) exhibited decreased IFNγ and GZMB expression compared to the CD8^+^ T cells incubated with BMDM from *Cx3cr1* null mice evaluated under the same conditions (**Figure 4I-J**). These results suggest that myeloid cell intrinsic E2/ER increases CX3CR1 expression leading to a suppression of CD8^+^ T cell activation. Thus, we conclude that E2 signaling attenuates type I IFN signaling in myeloid cells and suppresses T cell function in a CX3CR1-dependent manner.

#### Small molecule inhibitor of CX3CR1 reverse the immunosuppressive actions of E2 in the tumor microenvironment

To explore the translational importance of our findings, which linked CX3CR1 to ER-biology in macrophages, we evaluated the extent to which AZD8797, a small-molecule allosteric-inhibitor of CX3CR1 recapitulated the phenotypes observed upon genetic ablation of this receptor. Importantly, pharmacological inhibition of CX3CR1 suppressed E2-mediated tumor growth in preclinical models of NSCLC (LLC1), breast cancer (A7C11), and melanoma (B16F10) (**Figure 5A-C**). Comparative analysis of the intratumoral immune cells from placebo and drug-treated animals revealed an increase in MHC-class II expression in TAMs in the AZD8797 treated cells implying enhanced antigen-presenting capacity and pro-inflammatory functions. These changes coincided with increased CD8^+^ T cell activation (CD44/CD69^+^, GZMB^+^, IFNγ^+^, PD1^+^) (**Figure 5D- H**), reinforcing the conclusion that CX3CR1 is a mediator of E2-induced immunosuppression and highlighting the potential utility of this signaling axis as a therapeutic target in solid tumors.

**Figure 5:**
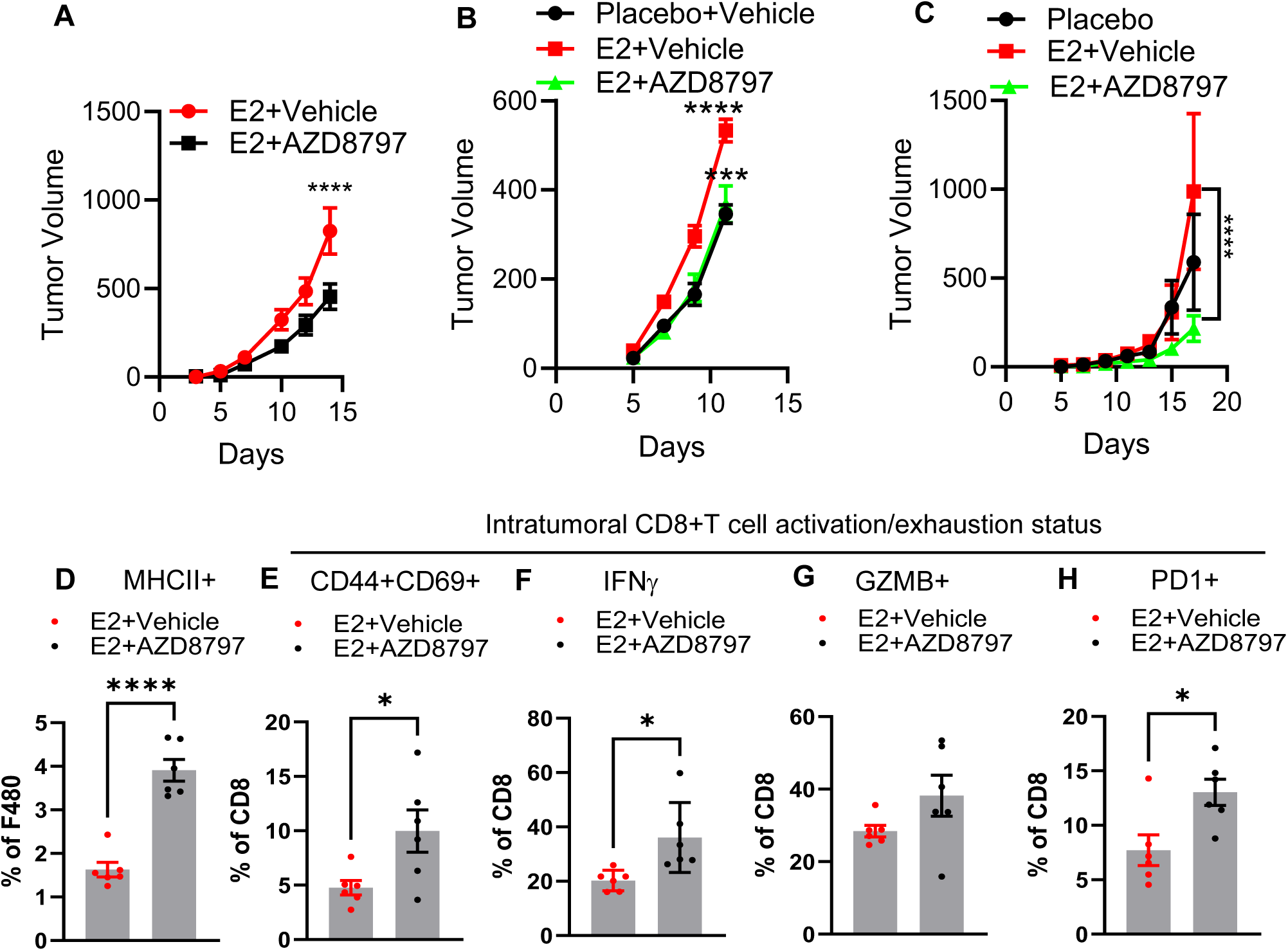
Small molecule inhibitor of CX3CR1 reverse the immunosuppressive actions of E2 in the tumor microenvironment. Syngeneic tumor growth of LLC1 (NSCLC) (**A**), A7C11 (breast carcinoma) (**B**) and B16F10 (melanoma) (**C**) injected in ovariectomized C57BL6/J mouse treated with E2 in presence or in the absence of CX3CR1i AZD8797 (A-C). Flow cytometry quantification of intratumoral M1 macrophages (MHCII+) (**D**) and CD44+CD69+ (**E**) IFNγ+(**F**) GZMB+ (**G**) and PD1+ (**H**) CD8+ T cells from experiment in 4B. Data represent mean ± S.E.M. Significance is calculated by Students t- test, two-way ANOVA followed by Bonferronni’s multiple corrections. *p<0.05, **p<0.01, ***p<0.001, ****p<0.0001.

### E2 increases NFAT activation to upregulate CX3CR1 expression in tumor-associated myeloid cells

Macrophages undergo rapid changes in chromatin dynamics and transcriptional reprogramming after efferocytosis of apoptotic tumor cells [42]. Thus, as an approach to identify the mechanisms by which ER/E2 impacts chromatin accessibility in myeloid cells and the activity of transcription factors that mediate these responses, we performed ATAC sequencing of chromatin isolated from LLC1 associated TAMs from ovariectomized mice treated ± E2. Genomic annotation of differentially accessible sites revealed enrichment of E2-responsive accessible sites in different regulatory regions of the genome. Promoter proximal regions comprised 25% of these sites, while 45% were at genic and 29% observed in distal intergenic sites (**Supplemental Figure 5A**). Cross-referencing differentially expressed vs accessible genes at promoter-proximal areas (5Kb upstream to 1Kb downstream of TSS) demonstrated high concordance between genomic accessibility and transcript expression (**Figure 6A, Supplemental Figure 5B-C**). Quantification of predicted transcription factor binding from ATAC-seq and their expression derived from RNAseq using DiffTF [43] revealed that E2 treatment enhanced the chromatin accessibility of predicted binding sites for multiple Ca^2+^ regulated transcription factors (NFATC1-C5, ATF3, CREB, EGR1, etc.), while decreasing accessibility of genes with binding sites for predicted IFN responsive transcription factors (IRF1, IRF8, IRF2) (**Figure 6B)**.

**Figure 6.**
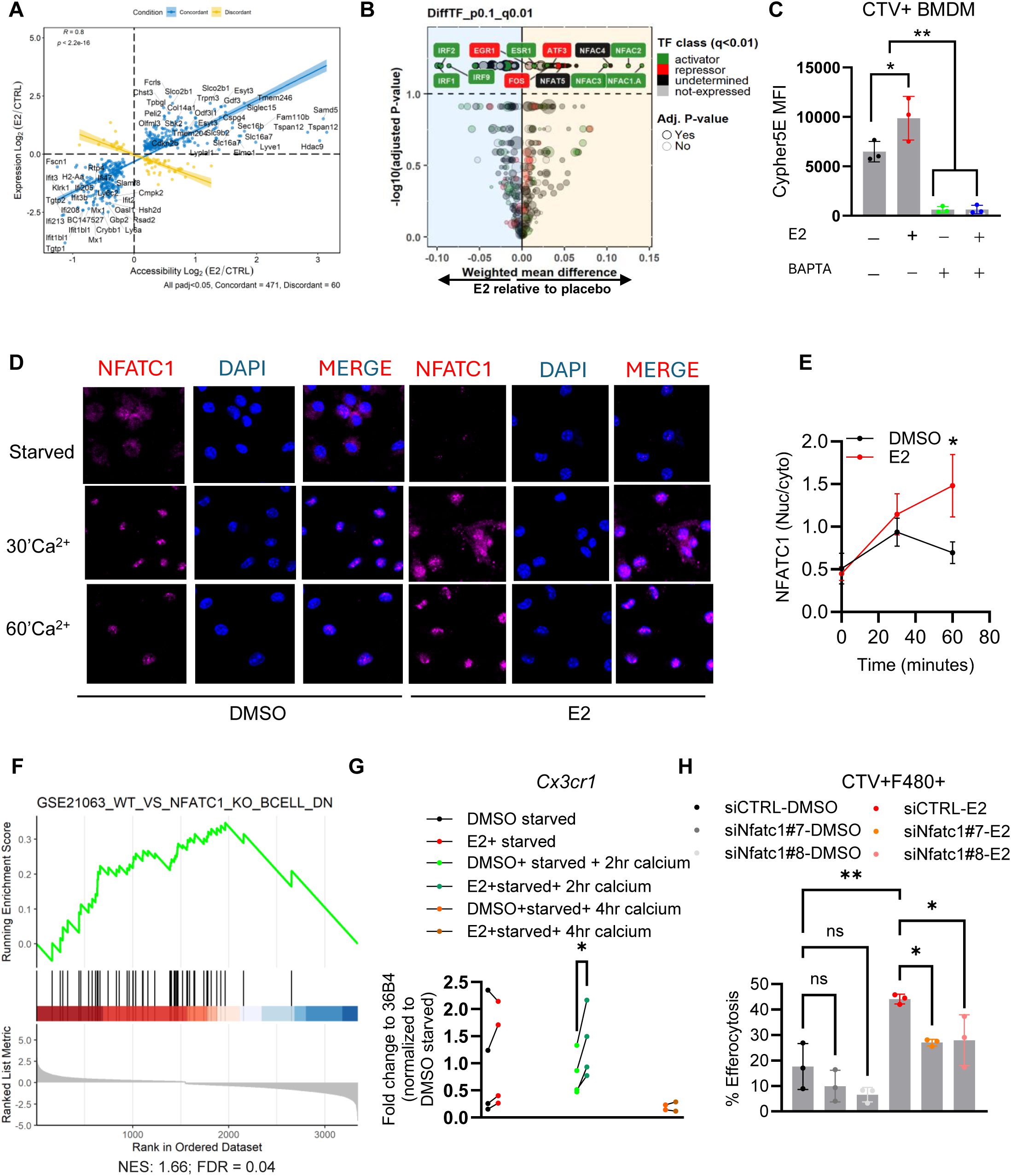
E2 increases NFAT activation to upregulate CX3CR1 expression in tumor-associated myeloid cells. Concordance between genes whose expression and chromatin accessibility were significantly changed (both up and down-regulated, *p<0.05) upon E2 treatment in intratumoral myeloid cells (n=4) (**A**). Predicted transcription factor binding sites in accessible chromatin from placebo and E2 treated intratumoral (LLC1) myeloid cells; red=predicted TF that acts as a repressor and its expression is decreased (E2 vs placebo) in RNA seq; green=predicted TF that acts as an activator and whose expression is increased in RNA-seq); black predicted TF that can act as both activator or repressor and whose expression is unchanged (**B**). Quantification of efferocytosis rate as measured by Cypher 5E staining in BMDM (n=3) treated ± E2 in the presence or absence of calcium chelator BAPTA (**C**). Immunofluorescence staining of NFATC1 in BMDM treated with TNFα (72 hrs) ± E2 for 48 hrs followed by starvation in Ca^2+^ free media for 4 hrs. Following 4 hrs starvation the cells were either left in starvation or treated with media containing 2mM Cacl2 for 30 minutes or 60 minutes. (**D**). Quantification of the ratio of mean fluorescent intensity of nuclear vs cytoplasmic NFATC1 from experiment 5D (n=3) (**E**). Gene set enrichment plot of NFATC1 target genes (GSE21063: WT vs NFATC1 KO B cells down) in intratumoral myeloid cells upon E2 treatment (E). Quantification of RNA expression of *Cx3cr1* from BMDM (n=2-4) treated with TNFa (72 hrs) ± E2 (48 hrs) followed by starvation (4hrs) and calcium addback for 2 hrs and 4 hrs (**G**). Percent efferocytosis of BMDM (n=3 biological replicates) that were transfected with either scrambled siRNA or siRNA-targeting NFATC1, followed by either DMSO or E2 treatment. Cells were co-treated with apoptotic CTV+ Jurkat cells in a 1:1 ratio for 45 mins following which their engulfment capacity was measured (**H**). Error bars represent± S.E.M. Significance was calculated by either Student’s t-test (G) one-way ANOVA followed by Tuckey’s multiple comparison (C), Sidak’s multiple comparison (H) or two-way ANOVA followed by Sidak’s multiple correction (E), *p<0.05, **p<0.01, **p<0.001, ****p<0.0001.

The observation that E2 exposure regulates the activity of calcium-regulated transcription factors was intriguing, as previous studies have reported a causal role for calcium in regulating multiple steps in the efferocytic cascade, specifically in phagosome maturation required for cargo degradation [44–47]. Indeed, we determined that pretreatment of macrophages with the intracellular calcium-chelator BAPTA AM inhibited both basal and E2 stimulated increases in apoptotic cargo degradation following efferocytosis as evidenced by the noted decrease in phagolysosomal acidification (decrease in cypher5HE staining) in CTV+ macrophages (**Figure 6C**). These findings implicate the NFAT family of transcription factors as mediators of the effects of E2 on efferocytosis as (a) their activity is dependent on the mobilization of intracellular calcium and (b) the binding sites for these transcription factors were among the most enriched in chromatin isolated from E2 treated myeloid cells. That NFATC1 directly regulates CX3CR1 expression in myeloid cells has been established by others [48]. Thus, we probed whether the increased expression of CX3CR1 in E2-treated macrophages could be attributed to changes in NFAT activity. Under unstimulated conditions, phosphorylated NFAT remains in the cytoplasm in an inactive state. Upon calcium stimulation, the phosphatase calcineurin dephosphorylates NFATC1, which results in NFATC1 translocation to the nucleus, where it promotes transcription of its target genes [49]. NFATC1 expression is low in macrophages at baseline but can be stimulated upon TNFα treatment [50]. To determine if E2 treatment impacts the nuclear translocation of NFATC1, TNF-exposed macrophages were treated ± E2, and NFATC1 activity was determined under both calcium starved and calcium-replete conditions (4hrs starvation followed by 30 or 60 mins calcium stimulation). The degree of nuclear translocation of NFATC1 was similar in placebo and E2/Ca^2+^ treated cells after 30 min. However, at later time points (60 mins), E2-treatment favored nuclear retention of NFATC1 compared to placebo control (**Figure 6D-E**). This result mirrored the results of studies performed *in vivo* where we determined that an established NFATC1 gene signature (GSE 21063) was enriched in RNA from E2-treated myeloid cell samples (**Figure 6F**). We extended these studies to show that the expression of *Cx3cr1*, was increased in TNF**α**-treated macrophages co-treated with E2 and that this activity required calcium (**Figure 6G**). We next interrogated the role of NFATC1 in E2-mediated efferocytosis. To this end, differentiated BMDMs were transfected with scrambled siRNA or siRNA targeting *Nfatc1* mRNA (**Supplemental Figure 5E**). BMDMs were further treated with either DMSO or E2 and the impact of these manipulations on efferocytosis was determined. While depletion of NFATC1 resulted in a partial reduction of efferocytosis in DMSO-treated macrophages the effect of this knockdown had a substantially greater effect on the efferocytotic capacity of E2-treated macrophages. We concluded from these experiments that the ability of E2 to increase efferocytosis efferocytic is mediated in large part through NFATC1 (**Figure 6H**).

### ER modulators enhance the efficacy of radiotherapy by suppressing efferocytosis

Our work has established a causal role for the E2/ER/CX3CR1 axis in promoting efferocytosis-mediated suppression of type I IFN signaling and, consequently, CD8^+^ T cell activation in the tumor microenvironment. As macrophages employ efferocytosis to remove apoptotic cancer cells from the tumor microenvironment, we hypothesized that inhibition of efferocytosis may improve the efficacy of interventions that result in an increase in intratumoral apoptotic cells (i.e. chemotherapies and radiation therapy). Previous studies have shown that debulking of tumors using ionizing radiation (IR) produces apoptotic cells as a byproduct [51, 52] and in preclinical models, pharmacological inhibition of targets that mediate apoptotic cell recognition by macrophages (Anti-PtDser and anti-MERTK) enhance IR efficacy [15, 53]. Thus, we hypothesized that inhibition of ER signaling, or its downstream targets, may enhance the efficacy of IR by attenuating efferocytosis and the associated suppressive effects of E2 on IFN signaling. Since ionizing radiation is used as a first-line treatment for NSCLC, we used the LLC1 model to test this hypothesis. In this model E2 treatment attenuated the effect of ionizing radiation (single dose, 5Gy) compared to its placebo-treated control (**Supplemental Figure 6A**). That monocyte migration and infiltration is initiated in response to radiation-induced necrosis in cancer cells has been reported by others [54–56]. Similarly, we demonstrated that radiation following E2 treatment was associated with increased Ly6C^+^ monocytes within the systemic circulation and a corresponding increase in Cx3CR1^+^ monocytes and macrophages within the tumor microenvironment (**Supplemental Figure 6B-D**). Given our observation that activation of the E2/CX3CR1 axis promotes efferocytosis-stimulated tumor growth, we sought to determine if inhibiting E2-ER signaling impacted the efficacy of radiation therapy. To this end, ovariectomized C57BL/6 mice were treated with and without radiation (5Gy), and the impact of E2 and a SERD (fulvestrant) on radiation efficacy were assessed. Tumor growth was not significantly impacted by low dose (5Gy) radiation (Placebo+Radiation vs Placebo+Veh), but the treatment of E2 (E2+ Radiation) facilitated tumor growth (**Figure 7A-B**). Importantly, the addition of fulvestrant (E2+ful+Radiation) reversed the inhibitory effects of E2 on IR suppressing tumor growth below that which was observed in Placebo+Radiation treated animals (**Figure 7B, Supplemental Figure 6E-G**). The number of CD206^+^ and CX3CR1^+^ macrophages were increased in the E2+IR treated samples compared to those subjected to IR alone, an activity that was significantly reversed upon the addition of fulvestrant (**Fig 7C-D**). The engulfment capacity of TAMs also showed a significant increase when E2 was combined with IR and a trend toward reversal was also noted with fulvestrant co-treatment (**Figure 7E**). These changes in macrophage phenotypes were also associated with an increased accumulation of activated IFNγ^+^ CD8^+^ and CD4^+^ and PD1^+^ CD8^+^ T cells (**Figure 7F-H**), consistent with the hypothesis that inhibition of estrogen signaling increases cancer immune surveillance to enhance IR efficacy.

**Figure. 7.**
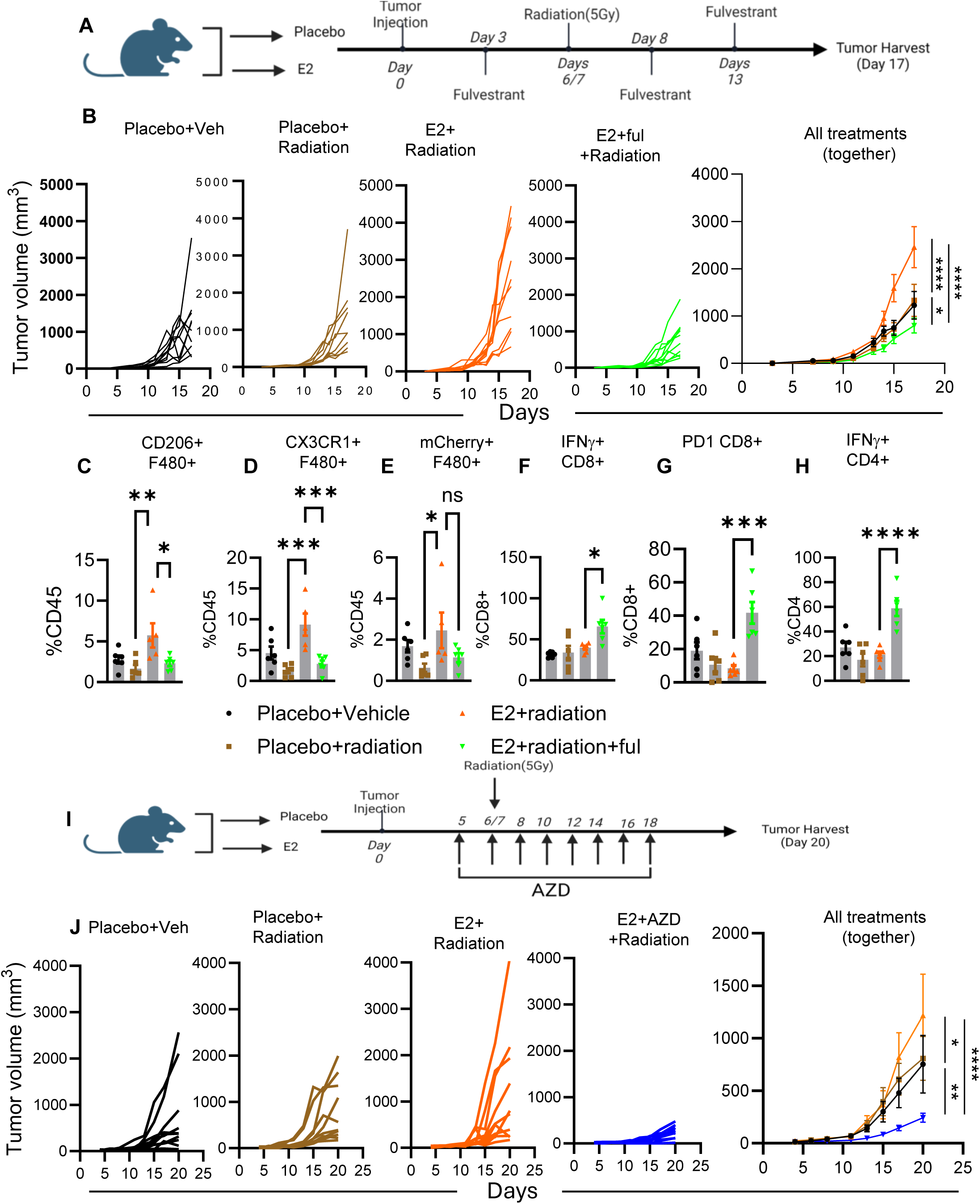
ER modulators enhance the efficacy of radiotherapy by suppressing efferocytosis. Schematic of combination treatment using radiation (5Gy) and SERD fulvestrant (A). Syngeneic tumor growth of LLC1 cells in ovariectomized mice treated with placebo, placebo+5Gy radiation, E2 + 5Gy radiation and E2+radiation+fulvestrant (25mg/kg) (n=10mice/group) (**B**). Flow cytometry quantification of CD206+ macrophages (**C**); CX3CR1+ macrophages (**D**); mCherry+ macrophages (**E**), IFNg+ CD8+ T cells (**F**); PD1+ CD8+T cells (**G**) and IFNg+CD4+ T cells (**H**) from tumors isolated from experiment described in 6B. Schematic of combination treatment using radiation and anti-CX3CR1i AZD8797 (**I**) Syngeneic tumor growth of LLC1 cells in ovariectomized mice treated with placebo; placebo+radiation (5Gy); E2+radiation (5Gy); E2+radiation+CX3CR1i (2mg/kg) n=10 animal/group (**J**). Data represents S.E.M. Significance was calculated by two-way ANOVA and pairwise comparison followed by Dunnett’s multiple corrections (B and J), one-way ANOVA and pairwise comparison followed by Dunnett’s multiple corrections (C, D, F, G and H), pairwise comparison followed by Sidak’s multiple comparisons (E) and (J). *p<0.05, **p<0.01, ***p<0.001, ****p<0.0001). Schematics in A and I were created by biorender.com.

We next assessed the impact of directly inhibiting CX3CR1 on IR efficacy. Ovariectomized mice injected with LLC1 cells were treated with or without E2 and with and without the CX3CR1 inhibitor AZD8797 in the presence or absence of IR. Studies of tumor growth revealed significant growth inhibition upon cotreatment with the CX3CR1 inhibitor and IR compared to IR alone (E2+IR and E2+IR+CX3CR1i) (**Figure 7I-J and Supplemental Figure 6H-I**). Taken together, these findings highlight the therapeutic potential of inhibiting the E2-ER-CX3CR1 signaling pathway as an approach to increase the antitumor efficacy and immune stimulatory impact of IR.

## Discussion

This study provides insights as to the mechanism by which the steroid hormone E2 influences the immune-modulatory functions of TAMs. We have previously demonstrated that E2 polarizes TAMs towards an anti-inflammatory state that limits CD8^+^ T cell activity and ICB efficacy [23]. Here we demonstrate that E2 promotes efferocytosis by TAMs by enhancing the expression of the macrophage intrinsic “find me” receptor CX3CR1 leading to a suppression of downstream IFN signaling. The induction of efferocytosis through CX3CR1 activation culminates in the suppression of pro-inflammatory IFN signaling in myeloid cells and subsequently with CD8^+^ T cell suppression and tumor progression. Importantly these immunosuppressive activities of E2 can be inhibited using a SERD or with a CX3CR1 antagonist (**Figure 8**). The availability of SERDs for clinical use suggests that our findings can be exploited in the near term to improve tumor immunity [57].

**Figure 8:**
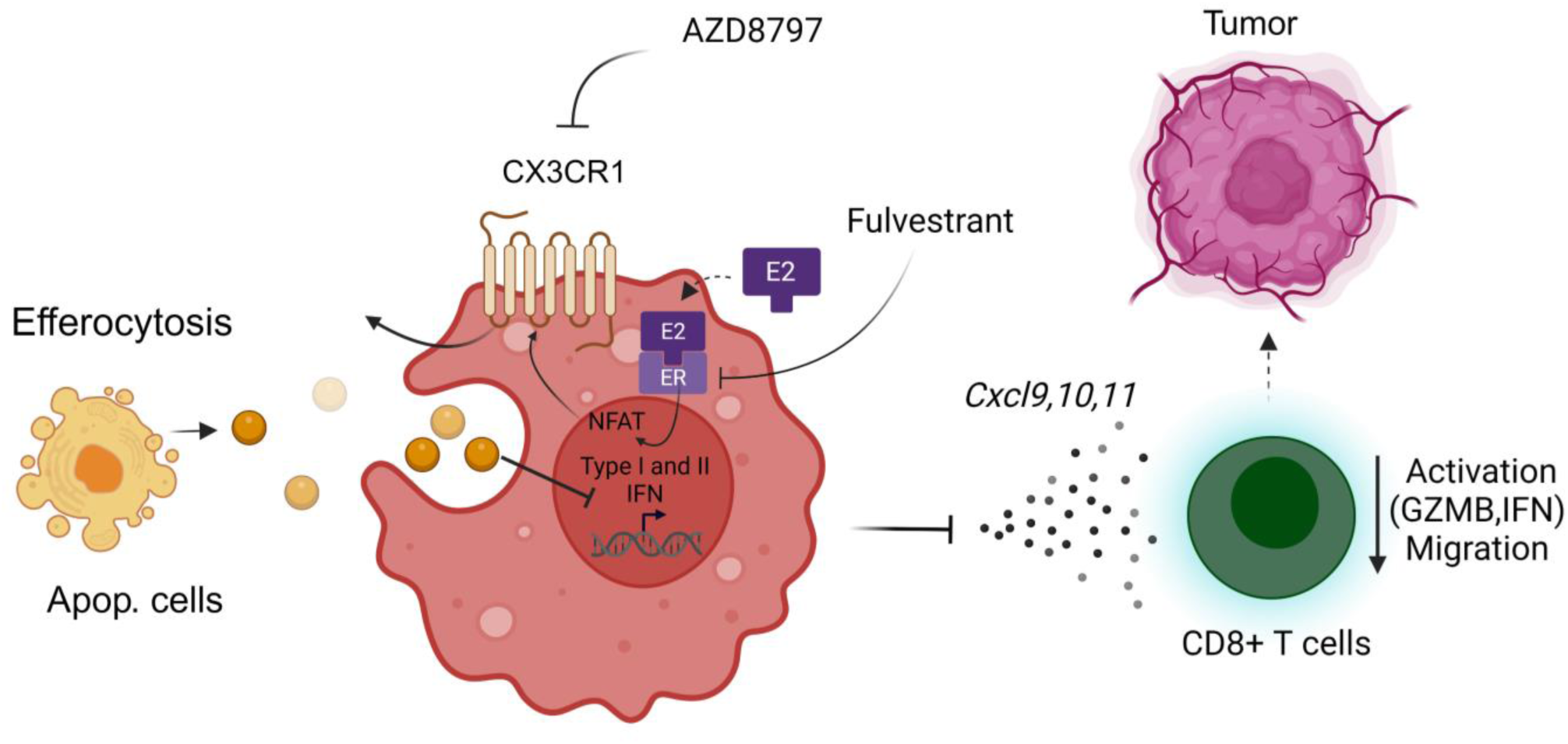
Schematic illustration of E2-regulated efferocytosis within the tumor microenvironment. E2 enhances the activity of transcription factor NFATC1 leading to increased expression of its downstream target CX3CR1 that enhances TAM efferocytosis and subsequent suppression of myeloid cells intrinsic type I IFN signaling.

While our understanding of macrophage biology has evolved over the last decade, the drivers of macrophage phenotypes and functionality that control their wound-healing and anti-inflammatory activities remain enigmatic. Macrophages are functional phagocytes that engage in the removal of dead/dying cells from tissues and are thus generally immune-suppressive. In normal physiology, the process of efferocytosis is required to prevent excessive inflammation at wound sites that may lead to cell and tissue damage and/or autoimmunity [9]. This may explain some of the positive effects of estrogens on wound healing and tissue repair [58]. Here we demonstrate that this evolutionarily conserved process enables estrogens to accomplish local immune suppression, allowing cancer cells to grow in an environment with limited immune surveillance.

The process of efferocytosis is regulated by a coordinated series of steps that allows macrophages to first sense ‘find me’ signals released from dead cells that promote TAM chemotaxis to the tumor microenvironment. This is followed by the interaction of ligands (e.g. PtDser) expressed by dead cells with specific receptors on macrophages that interpret these “eat me” signals resulting in dead cell engulfment followed by destruction through autophagy or LC3- associated phagocytosis (LAP) [59–61]. Here, we show that E2 induces the expression of the ‘find me” receptor CX3CR1 whose activation is associated with macrophage polarization towards an immune-suppressive state [62]. Previous studies have demonstrated that CX3CR1 activation in macrophages induces JAK2/STAT3 signaling while suppressing NFκB activation [63, 64]. Interestingly, activation of the JAK2/STAT3 axis or suppression of NFκB pathway has also been shown to be associated with suppression of proinflammatory type I IFN signaling [65]. In line with these observations, we demonstrated that E2/ER signaling controls the engulfment capacity of CX3CR1^+^ macrophages in the tumor microenvironment and that deletion of CX3CR1 reverses the tumor growth-promoting effects of E2 and its suppressive effects on type I IFN signaling. While we cannot exclude roles for CX3CR1 signaling in other immune cell types, e.g. T cells, as being involved in E2 biology in tumors, our work highlights the primacy of CX3CR1 in macrophages in this biology. Indeed, tumor cells admixed with CX3CR1^-/-^ BMDM suppressed the tumor growth phenotype compared to those that were mixed with CX3CR1-het BMDM in an E2-exposed tumor microenvironment. Thus, while other studies have reported that T cell-intrinsic CX3CR1 signaling/expression predicts ICB efficacy, T cell memory, and immunosurveillance, we did not observe any detrimental changes to tumor-infiltrating lymphocytes upon CX3CR1 deletion [66, 67]. This could be due to the nature of the microenvironment established in the tumor models we used as these which are dominated by the presence of myeloid cells and have limited T cell infiltration. Moreover, *in vitro,* we demonstrated that CX3CR1 deficiency/inhibition in BMDMs reversed E2/apoptotic cell-enhanced, macrophage-dependent suppression of CD8^+^ T cells. This establishes CX3CR1 as a key downstream effector of E2 that promotes macrophage-mediated immunosuppression within the tumor microenvironment.

ATAC sequencing analysis of chromatin from TAMs isolated from E2-treated tumors revealed increased accessibility of regions that are enriched for canonical DNA binding sites of calcium-regulated transcription factors. This was interesting given the established roles of calcium in multiple steps in the efferocytotic cascade [44–47, 68]. We demonstrated that E2-dependent calcium mobilization promotes activation of NFATC1 to enhance the transcription and expression of CX3CR1 within TAMs. While the role(s) of NFATC1 in T cell activation is well established, little is known about the function of this transcription factor in macrophages [50, 69–71]. It is significant therefore that we have shown that by activating a transcriptional program that facilitates macrophage efferocytosis, NFATC1 promotes macrophage polarization towards an anti-inflammatory state. While we established a role for calcium and calcium-associated transcription factors in the regulation of efferocytosis in E2-treated macrophages, the underlying mechanism(s) by which this hormone increases intracellular calcium remain to be determined. In previous work, performed in breast cancer cells, we have shown that estrogens facilitate the release of calcium from intracellular stores [72]. Whether or not a similar pathway exists in macrophages is currently under investigation. Regardless, the results of our studies suggest the potential utility of targeting the estrogen signaling axis as a means to regulate efferocytosis in cancer and in other diseases where its dysregulation is associated with specific pathobiology [73–77].

The process of efferocytosis enables the rapid clearing of apoptotic cells via LC3-dependent phagocytosis or by autophagy [35]. These processes are equally important for apoptotic cargo clearance as the accumulation of apoptotic corpses within macrophages can result in dsDNA and RNA-mediated activation of the dsDNA/RNA sensor STING/RIG-I/MDA5, and induction of inflammatory signaling. In normal physiology, these responses are desirable and needed to limit inflammatory responses in wounds that can lead to tissue damage. In tumors, however, efferocytosis leads to immunosuppression and favors disease progression. Indeed, we have demonstrated that E2-treated macrophages within the tumor microenvironment suppress the activation of type I IFN signaling, which has been shown to limit processes such as antigen presentation, MHCI and II expression, proinflammatory cytokine release, and the release of chemokines that promote T cell migration, activation and cytotoxicity [78]. Multiple studies have demonstrated that injection of low doses of type I IFN into tumors can promote T cell-dependent anti-tumor activity, findings that led to the clinical development of IFNα-2b for the treatment of metastatic melanoma, chronic myelogenous leukemia and hepatocellular carcinoma. Despite showing encouraging results in patients, the use of IFNα-2b in this manner is limited by side effects arising from its ability to induce excessive non-specific immune stimulation [79]. Our preclinical results show that clinically available ER-directed endocrine therapies can reverse E2-mediated suppression of myeloid cell-intrinsic type I IFN signaling and, in doing so, enhance adaptive immune responses. Thus, endocrine therapies may serve as a useful alternative to strategies that use IFNα-2b and/or other IFN inducing agents (e.g., STING agonists or virotherapy) to enhance the activity of IFN signaling within the tumor microenvironment. The restricted expression of ER makes this approach particularly appealing. Further, our studies have indicated that CX3CR1 is a key downstream mediator of the inhibitory actions of the ER/E2 on IFN signaling and thus drugs which target this receptor may be a useful approach to increase tumor immunity.

It is widely recognized that radiation therapy exerts its therapeutic effects in part by inducing apoptosis-mediated cell death [80, 81]. Importantly, targeting components of the efferocytosis machinery or blocking ligands (PtDser) that assist macrophages in recognizing apoptotic cells can enhance radiotherapy efficacy [15, 21, 82]. In our preclinical mouse models of non-small cell lung cancer (NSCLC), we discovered that the inhibition of ER-signaling with clinically relevant endocrine therapies enhances the activity of radiation therapy by inhibiting efferocytosis and activating adaptive immunity. Given that the current standard of care for locally advanced NSCLC is radiotherapy followed by immune checkpoint blockade (ICB), there is considerable interest in developing radiosensitizers to enhance immune activity in a post-irradiated tumor microenvironment to increase the efficacy of ICB [83]. Our studies thus highlight the potential of using neoadjuvant endocrine therapy as a route to improve antitumor immune responses to IR and ICBs.

In conclusion, we have defined a novel ER/NFAT/CX3CR1 axis that controls phagocytic clearance of dead cells by tumor-associated macrophages thereby suppressing anti-tumor immunity and limiting the efficacy of radiation therapy. The ability of ER-directed endocrine therapies and or CX3CR1 inhibitors to reverse macrophage-mediated immune-suppression in post-irradiated tumors provides a compelling rationale for the clinical development of combination therapeutic approaches that target this axis to enhance RT responses, and potentially other modalities in patients with cancer.

## Methods

### Mice

All experiments were performed according to guidelines listed in IACUC, Duke University. C57BL/6J (Catalog #000664), Lysozyme-MCre (B6.129P2-*Lyz2tm1(Cre)Ifo*/J (Catalog #004781) [84], Mx1-gfp (B6.Cg-*Mx1^tm1.1Agsa^*/J) (Catalog #033219) [37], CX3CR1^gfp/gfp^ B6.129P2(Cg)- *Cx3cr1^tm1Litt^*/J (Catalog #005582) [41] were purchased from The Jackson Laboratory, Maine USA. Age-matched mice were used for all the studies described. LysMCre animals were further bred with *Esr1^fl/fl^* mice (a gift from Ken Korach, National Institute of Environmental Health Sciences [NIEHS], NIH, Durham, North Carolina, USA) to generate *Esr1fl/fl* LysMCre and littermate control LysMCre and *Esr1^fl/fl^* mice. CX3CR1^gfp/gfp^ mouse were bred with WT-C57BL/6J mouse to generate Cx3CR1+/gfp. These animals were further crossed among themselves to generate experimental cohorts of genotype Cx3CR1^+/gfp^ and CX3CR1^gfp/gfp^. Mx1-gfp were bred among themselves to maintain a homozygous Mx1^gfp/gfp^ genotype. The mice were housed in secure animal facility cages on a 12-hour light/12-hour dark cycle at a temperature of approximately 25°C and 70% humidity. Mice had ad libitum access to food and water.

### Cell lines, cultures and reagents

RAW264.7, J774A, B16F10, LLC1 and Jurkat cells were purchased from American Type Cell culture Association. LLC cells expressing mCherry-Spectrin were a kind gift from Dr. Douglas Green’s lab at St. Jude Meical Center. A7C11 cells were kindly provided by Dr. Jose.R Conejo Garcia, Duke University. BPD6 cells were a kind gift from Dr. Brent Hanks, Duke University. RAW264.7 and J774A, B16F10 cells were cultured in Dulbecco’s Modified Eagle’s Media (DMEM) media supplemented with 10% fetal bovine serum (FBS), 1% non-essential amino acids (NEAA) and sodium pyruvate (1mM). LLC1 and LLC-mCherry-Spectrin cells were cultured in DMEM supplemented with 10% FBS. Jurkat cells were cultured in RPMI 1640 medium supplemented with 10% FBS and 1mM- β-mercaptoethanol. A7C11 cells were grown in RPMI 1640 medium supplemented with 10% FBS, 1% Na-Pyruvate, 1% NEAA and 1 2- Mercaptoethanol. Cells were cultured in a 37°C, 5% CO2 incubator and were split using 0.25% Trypsin 2-3X/weeks (1:10 ratio). The non-adherent Jurkat cells were sub-cultured in 1:10 ratio by spinning down cells + supernatant at 1500rpm for 5 minutes.

### Surgical Ovariectomy

7-8 weeks old C57BL/6J female mice were used for ovariectomy. Mice were anesthetized in an inhalation chamber containing 2% Isoflurane and maintained in 50% of the dose of isoflurane (1%) via nose cone throughout the entire surgical procedure. Prior to surgery, mice were injected subcutaneously with carprofen (5mg/kg). The area below the ribs was shaved with an electronic razor. Sterilization of skin was performed by rubbing with betadine and alcohol (3X alternating). Just above the ovary fat pad a horizontal incision was made through the skin, followed by a vertical incision through the abdominal muscle wall. The ovary was then externalized and surgically removed using a cauterizing scissor. Fat pads were replaced at original position, and muscle walls were opposed and sutured (1-2 stitches). Following the suturing a single drop of bupivacaine (0.25%) was added on top of the incision site. The skin was opposed followed by an administration of wound clip at the incision site. Similar procedure was repeated for the other ovary. The mouse was then removed from anesthesia and kept in a sterile clean cage and monitored continuously until they regain consciousness.

### In vivo Tumor Growth

For subcutaneous tumor models B16F10 (0.5x10^5^), LLC1 (2.5 x10^5^) and LLC1-mCherry-Spectrin (2.5X10^5^) were either injected in the right flank or around the chest area. A7C11 cells (0.2x10^5^) were injected in the left second mammary fat pad. Tumors were allowed to grow, and tumor size was measured 3x/week. Tumor volumes were calculated by the formula T= Lx (WxW)/2. For tumor studies, animals were euthanized before or when tumors reached a maximum size of 2000mm3 as specified by IACUC, Duke University.

### Single Cell Suspensions of tumors

Animals were euthanized when the tumors reached a volume of approx. ≥ 2000mm3. The tumors were isolated and were minced into small pieces using mechanical shopping methods with sterilized razor blades or by using the C tubes (Miltenyi Biotec, Catalog # 130-093-237) and dissociating using gentleMACS dissociator (Miltenyi Biotec, Catalog Number: 130-093-235). The chopped tissues were dissolved in DMEM+5% FBS medium containing 100µg/ml DNase I (D5025, Sigma-Aldrich) and 1mg/ml collagenase (Collagenase A, Cat# 10103586001, Sigma Aldrich). The resulting cell suspensions were then filtered through a 40µm strainer to produce single cell suspensions. The enzymes were diluted by adding additional media then spun down to remove media. Red blood cells were lysed with the addition of ACK lysis buffer (Cat# A1049201, ThermoFisher Scientific) for 3-4 minutes at room temperature. Following red blood cell lysis cells were washed with PBS, counted before proceeding to flow cytometry staining or magnetic bead-based isolation.

### IFNAR Blocking Studies

For the purpose of IFNAR depletion, C57BL/6J mice were injected with 500µg/mouse of a mouse anti-IFNAR antibody (clone MAR1-5A3, cat # BE0241 BioXCell) or mouse IgG1 isotype control (cloneMOPC21, cat # BE0083,) diluted in sterile PBS, 24 hours before tumor injection and every 3 days after tumor injection.

### In vitro Bone Marrow Isolation and Differentiation

For this purpose, bone marrow cells were aseptically collected from female C57BL/6J (8-12 wks) mice by crushing the femurs and tibias in PBS, 1% FBS and 2mM EDTA. Cells were spun down and the pellet was mixed with ACK buffer to lyse the red blood cells for 2 minutes with intermediate vortexing. The ACK buffer was then diluted in PBS (1:10). The solution was filtered through a 40µm strainer to remove bone fragments. The filtrate was spun down and the supernatant was aspirated to remove the ACK buffer. To differentiate bone marrow cells to macrophages the cells were plated in Phenol Red Free-DMEM media supplemented with 10% heat-inactivated charcoal-stripped serum in the presence of 30 ng/ml MCSF (Cat# 312-02, PeproTech) and 1% Pen/Strep. Following 3 days of culture, cells were supplemented with 50% of fresh media. On day 6 the media was removed and replaced with fresh media. When the cells are fully differentiated to macrophages on day 7, treatments were initiated.

### Human Macrophage Differentiation and Culture

PBMC isolated from de-identified human donors were plated for an hour in the presence of DNAse following which the floating cells were removed. Phenol Red Free AIMV (A3830801, Thermo Fisher Scientific) media was supplemented in the presence of 50ng/ml of mCSF (Catalog #78057.2, Stem Cell Technologies). Media was supplemented on day 5 and fully differentiated macrophages were obtained on day 7.

### In vitro Drug Treatments

Macrophage cultures (BMDM and hMDM) were treated with DMSO and 1nM 17β-E2 for 48 hrs while J774 and RAW 264.7 cells were treated with DMSO and E2 for 72 hrs. J774A cells were treated with Cytochalasin D (1nM) for 16 hrs. BAPTA (10µM) was added to the BMDM 1hr prior to start of the efferocytosis assay. Ionomycin was treated at 500nM for 1hr.

### In vivo Drug Treatments

Mice were injected with the SERD fulvestrant (Cat# HY-13636 MedChemExpress) 48hrs following tumor injection at a dose of 25mg/kg via intramuscular route. Following the initial injection fulvestrant treatment was administered to the mice every 5 days. Corn oil (Cat # C0136, Spectrum Chemical MFG Corp) was used as vehicle for fulvestrant which was administered at the same frequency (every 5 days/intramuscular route) to all other animals. For the CX3CR1 inhibitor studies, mice were injected with CX3CR1i AZD8797 72 hours following tumor injection at 2mg/kg via intraperitoneal routes. Beta Cyclodextran (βP-HCD) (Cat # 89428- 750, Sigma Aldrich) was used as vehicle which was administered to all other animals. Following the initial injection AZD8797 was administered to the animals every 2 days till the end of the study.

### siRNA-mediated knockdown

Differentiated macrophages were transfected with either non-targeting control or siRNA targeting NFATC1 (30pmoles, Horizon Discovery) with Lipofectamine RNAiMax (Cat# STEM00003, Thermo Fisher Scientific) following the manufacturer’s instruction. Media was changed 24 hrs later and the cells were collected for downstream analysis 48 hrs after transfection. The sequences of siRNA are listed in supplementary **Table I**.

### Flow Cytometry and Cell Sorting

Single cells suspensions of excised tumors (10^6^ cells in 50µl) or in vitro macrophage cultures were incubated with Live/dead fixable dead cell stain in PBS for 10 mins at 4°C. Cells were spun down at 1000xg and were incubated with anti-CD16/32 (Catalog# 14-0161-85, ThermoFisher Scientific) in flow buffer (10gms BSA in 1L PBS) for 15 mins. Following this, cells were stained with an antibody cocktail in Brilliant Stain Buffer (Cat# 566349, ThermoFisher Scientific) for 30 minutes at 4 degrees. The antibodies used are listed in Supplementary Table II. For intracellular staining cells were fixed and permeabilized using the eBioscience Foxp3 Transcription Factor Staining Buffer Set (Cat# 00-5523-00, ThermoFisher Scientific) followed by intracellular staining with the desired antibody overnight at 4°C. Following intracellular staining the samples were further fixed with BD cytofix solution. Multicolor flow cytometry was performed in BD Fortessa 16 color analyzer. FACS results were analyzed by FlowJo_V10 software (FlowJo, LLC). Sorting of tumor associated myeloid cells (CD45+ CD11b+ Live dead ^neg^, propidium Iodide (PI) ^neg^) were performed using Astrios EQ high speed sorters.

### In vitro Efferocytosis Assay

For the purpose of the efferocytosis assay, apoptosis was induced in LLC1- mCherry Spectrin cells with a combination of BCL2 (ABT737) (Cat# 6835, Cayman Chemicals) and MCL1 (S63845) (Cat# 21131, Sapphire North America) inhibitors as described previously [35]. Apoptosis was induced in Jurkat cells by applying UV (250mj/cm^2^) following which cells were placed in 5% CO2 incubator for 3 hrs to recover. For labelling of apoptotic Jurkat, cells were incubated with Cell Trace Violet (CTV violet) (5µM) and Cypeher5 E (8µM) for 20 mins in 37°C waterbath. Cells were then washed with PBS, counted and added to macrophages at desired ratios as indicated in each experiment. Macrophages were incubated with apoptotic cells for 45 minutes following which the cells were thoroughly washed with PBS and stained with F480 prior to flow cytometry.

### In vivo Efferocytosis Assay

For this purpose, 4x10^6^ apoptotic LLC1-mCherry-Spectrin cells were injected in the peritoneal cavity of the mice. The animals were euthanized after 1 hr and peritoneal cavity was flushed with 5 ml of PBS followed by collection of peritoneal lavage. The collected lavage was spun down, treated with ACK to lyse the red blood cells and then proceeded to stain with antibodies followed by flow cytometry to detect macrophages that are mCherry+.

### In vitro T cell Activation Assay

CD8+ T cells were isolated from the spleens of C57BL/6J or mice with magnetic bead-based CD8 T cell isolation kit (Cat # 19853, StemCell Technologies). T cells from naïve mice were stained with 5µM Cell Trace Violet (Cat# C34557, ThermoFisher Scientific) for 20 minutes in pre-warmed PBS following which the cells were washed twice with PBS. Stained T cells were then counted and plated in 96-well plates coated with anti-CD3 antibody (0.5µg/ml, Cat# 19851, ThermoFisher Scientific) anti-CD28 antibody (1µg/ml, Cat# 16- 0281-86, Thermo Fisher Scientific) at desired density (100,000 T cells/20,000 BMDM) in the presence of IL2 (50ng/ml) (Cat# 212-12, PeproTech). 72 hours following plating, cells were incubated with protein transport inhibitors brefeldin (Cat# 00- 4506-51, Thermo Fisher Scientific) and monensin (Cat#00-4505-51, Thermo Fisher Scientific) for 6 hours at a final concentration of 2µM monensin and 3µg/ml brefeldin following which they were collected and were processed for staining for flow cytometry.

### Ex-vivo Melanoma Tissue Culture

Freshly extracted melanoma tissue was sliced to size (2x2mm) and plated in 24 well plates in phenol-red free AIM V media supplemented with 5µg/ml insulin (GIBCO, Catalog#12585-014), 10ng/ml EGF (Catalog # 236-EG), 500ng/ml hydrocortisone (Sigma-Aldrich Catalog # H0135-1) and 1x Pen-Strep + fungizone (100 X Anti-Anti reagent, Invitrogen catalog number 15240-062). The tissues were treated with desired treatments for 48 hrs following which the media and tissues were separately collected for cytokine and RNA analysis.

RNA extraction and quantitative PCR: Tumor infiltrating myeloid cells were isolated from tumors either using a CD11b isolation kit (catalog # 18970, StemCell Technologies) or by cell sorting. RNA was isolated using RNA UCP Micro kit (catalog # 73934, Qiagen). RNA from BMDM were collected by Aurum total RNA isolation kit (BioRad Catalog # 7326820). Human melanoma tissues were collected and stored in RNA later. For extracting RNA, tissues were dissolved in TRIzol Reagent (Life Technologies, Catalog # 15596018) + chloroform (1:5) and spun at 14000rpm for 15 mins followed by the isolation of aqueous phase and isolating RNA using the RNAeasy mini kit (Qiagen, Catalog # 74014). cDNA synthesis was performed using the iScript cDNA synthesis kit (Cat# 170-7691). Quantitative amplification was performed using Sybr Green (Cat# 1725124, Bio-Rad) and the CFX-384 Real Time PCR detection system. Primers used are listed in Supplementary Table III.

### RNA-sequencing

Myeloid cells were isolated from LLC1 tumors of either placebo or E2-treated animals by fluorescent activated cell sorting using antibodies against CD45, live/dead, propidium iodide and CD11b. Live/dead^neg^ PI^neg^, CD45^+ive^, CD11b^+ive^ cells were sorted and RNA was extracted using Qiagen RNA micro kit according to manufacturer’s instruction. 100ng of total RNA was used for library preparation. Libraries were sequenced using 50bp paired-end reads on the Nova-seq 6000 platform (Illumina).

### ATAC sequencing

Myeloid cells were isolated from LLC1 tumors as described above. Assay for Transposable Element (ATAC-seq) was carried out using the Omni ATAC protocol [85]. Nuclear extraction was performed using 50,000 myeloid cells/sample. Briefly, cells were centrifuged for 1000xg for 15 minutes at 4 °C. The resulting pellet was resuspended in 50µl of RSB buffer (10mM Tris-HCl, pH 7.4, 10nM NaCl, 3mM MgCl2) containing 0.1% NP-40,1% Tween-20 and 0.01% Digitonin.

Cells were incubated on ice for 3 minutes following which the cells were resuspended again in 1ml RSB containing 0.1% Tween-20. Cells were spun down again following which a transposition reaction was carried out (37 °C for 30 minutes at 1000rpm mixing) using Tn5 transposase (Catalog # FC-121-1030) in the presence of 10% Tween-20 and 1% Digitonin. Immediately following the transposition reaction, DNA was purified using the Qiagen MinElute Kit (#28204). The DNA was further cleaned using Zymo DNA clean and concentrator-5 kit (Cat # D4014). Transposed DNA was then amplified using Nextera barcoded primer pair (Supplementary Table IV) with NEBNext High-Fidelity PCR master Mix (Catalog# M0541) according to manufacturer’s instructions. One-tenth of the reaction was used in a pilot qPCR reaction to determine optimal amplification cycles for library construction. The remaining nine-tenths of the samples were then subjected to the predetermined PCR amplification cycles using the same NEBNext High Fidelity PCR reaction. The Amplified Library was purified using 1:1 Ampure XP beads. Library quantification was performed using NEB Library Quantification kit (Catalog # E7630, New England Biolabs). The library was pooled and was sequenced on a Nova-seq 6000 sequencer (50 bp paired-end sequencing).

### Immunofluorescence

BMDM were differentiated on glass coverslips (VWR Catalog # 48380- 068) placed in a 6-well plate. Fully differentiated and treated BMDM were washed 3X with PBS and were fixed with 4% paraformaldehyde for 10 minutes at room temperature. The coverslips were again washed 3X with PBS following which the samples were blocked in PBS+0.1% Triton-100 +5% goat serum for 1 hr followed by incubating with NFATC1 antibody (1:100, Thermo Fisher Catalog # MA3-024) diluted in blocking buffer for 16 hrs. The slides were washed 3X with PBS followed by incubation with a secondary antibody, Alexa Fluor 594 tagged goat anti-mouse (1:500, Thermo Fisher Scientific, Catalog # A11032) diluted in blocking buffer.

The coverslips were washed with PBS 3X to remove non-specific antibody binding, DAPI (1ug/ml) was then added for 5 mins. Excess DAPI was washed, and the coverslips were mounted on glass slides with Vectashield Antifade Mounting Medium (Vector Laboratory, Catalog # H1000-10). Images were obtained in Zeiss 880 inverted confocal microscope using a 63X- oil objective. For each condition images were taken from 3 biological replicates (3 images/condition). Images were quantified by FIJI software.

### Radiation Study

Tumors were visualized using the X-ray system for imaging [fluoroscopy] & irradiation was performed using SARRP small animal radiation research platform (Xstrahl).

Anesthetized mice (w/ 2 L/min oxygen & 2 % isoflurane) were held on the specimen positioning stage and the mice were treated with a single dose of 5Gy fractions of focal, image-guided tumor radiation using the variable collimator (220 kVp, 13 mA X-rays using a 0.15 mm Cu filter). As mice are anesthetized, acclimation to the device before the radiation procedure was not needed. Additional brief use of physical restraint (surgical tape) was used on tail or extremities as an additional precaution and lasted less than 15 min. Mice breathing and distress are consistently observed through a camera located inside the machine.

### Bioinformatics Analysis

RNAseq reads were aligned with mm10 reference genome and gene counts were obtained using STAR aligner [86]. Differential expression analysis was performed using DESeq2 [87] with recommended settings. Enhanced volcano package (https://github.com/kevinblighe/EnhancedVolcano) was used to make the volcano plots.

Pathway enrichment analyses were done using ClusterProfiler [88]. ATACseq reads were processed using nf-core/atacseq (ver 1.1.0) pipeline (<u>10.5281/zenodo.2634132</u>). In brief, the reads were trimmed using Trim Galore (https://www.bioinformatics.babraham.ac.uk/projects/trim_galore/) followed by alignment to mm10 using BWA. Picard was used to mark duplicates (https://broadinstitute.github.io/picard/).

Bigwig files with normalized coverage tracks were generated from merged replicates scaled to 1 million mapped reads and read coverage over regions of interest were created using Deeptools [89]. Differential accessibility sites were determined by DESeq2 [87]. Differential transcription factor binding analysis was performed by DiffTF [43]. Genomic annotation of peaks was performed using Homer [90] and ChIPseeker. Data wrangling and plotting were done in R using ggpubr (https://rpkgs.datanovia.com/ggpubr/), Tidyverse (https://www.tidyverse.org/) and ggplot2 (Wickham, 2016).

### Statistics

Statistical analysis was performed with GraphPad Prism 8.0 (GraphPad Software), using either a 2-tailed Student’s *t* test or 1- or 2-way ANOVA. For both 1-way and 2-way ANOVAs, a post-test analysis was performed using Bonferroni’s or Tuckey’s multiple correction. The number of replicates is indicated in the figure legends. A *P* value of less than 0.05 was considered statistically significant.

## Acknowledgement

We thank Anna Cartaya Tellacheya from Light Microscopy Core Facility for helping with the microscopy studies and Nerissa Williams of Duke Radiation core for help with the radiation experiments. We acknowledge Kiana Gunn for her help with macrophage- apoptotic cells co-mixing experiment. We would also like to acknowledge the assistance of Duke Sequencing and Genomic Technology Shared Resource for generations of RNA sequencing and ATAC sequencing datasets and the DCI Flow Cytometry shared resources for assistance with cell sorting and flow cytometry equipment.

## Author Contributions

B.C, C.Y.C and D.P.M conceptualized and designed all the experiments. S.R.F and G.B provided reagents for the project. B.C, P.C, M.B and D.C performed majority of experiments and analysis with help from A.G, R.S, S.A, M.K, D.M, F.L and J.K.B. Animal experiments were supervised by B.C and S.W. D.C, A.R and P.K.J helped with collection and analysis of chart review studies. The manuscript was written by B.C and D.P.M. with critical input from C.Y.C, P.C, M.B, S.A and S.R.F. The project was supervised by D.P.M. Conflict of Interest: The authors declare no conflict of interest.

