## Supplementary Materials for "Estrogens increase cancer cell efferocytosis to establish an immunosuppressive tumor microenvironment"

**Supplemental Figure 1. E2 suppresses myeloid cell-intrinsic type I interferon signaling**

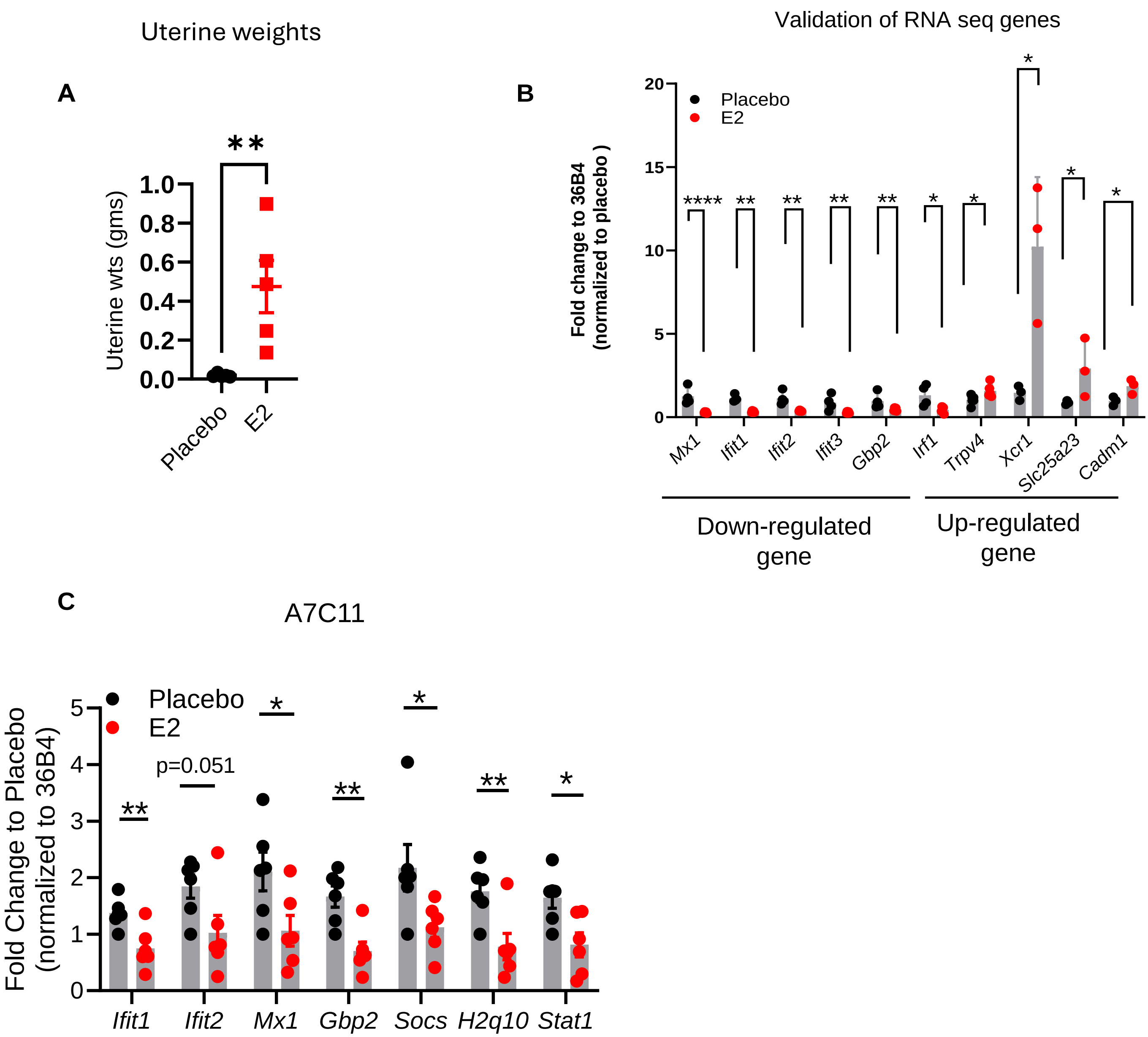


**Supplemental Figure 1**: **E2 suppresses myeloid cell-intrinsic type I interferon signaling. Related to Figure 1.** Uterine wet weights of ovariectomized animals treated ± E2 to determine E2 exposure (**A**). Validation of select up and downregulated genes from additional CD11b+ myeloid cells isolated from LLC1 tumors treated ±E2 (**B**). Expression of ISGs in myeloid cells isolated from A7C11 breast tumors ± E2 (n=6) (**C**). Data represent mean ± S.E.M. Significance is calculated by Students t-test, (*p<0.05, **p<0.01, ****p<0.0001.)

**Supplemental 2: Myeloid Cell Intrinsic Esr1 gene signature predicts survival in patients with varied solid tumor types**

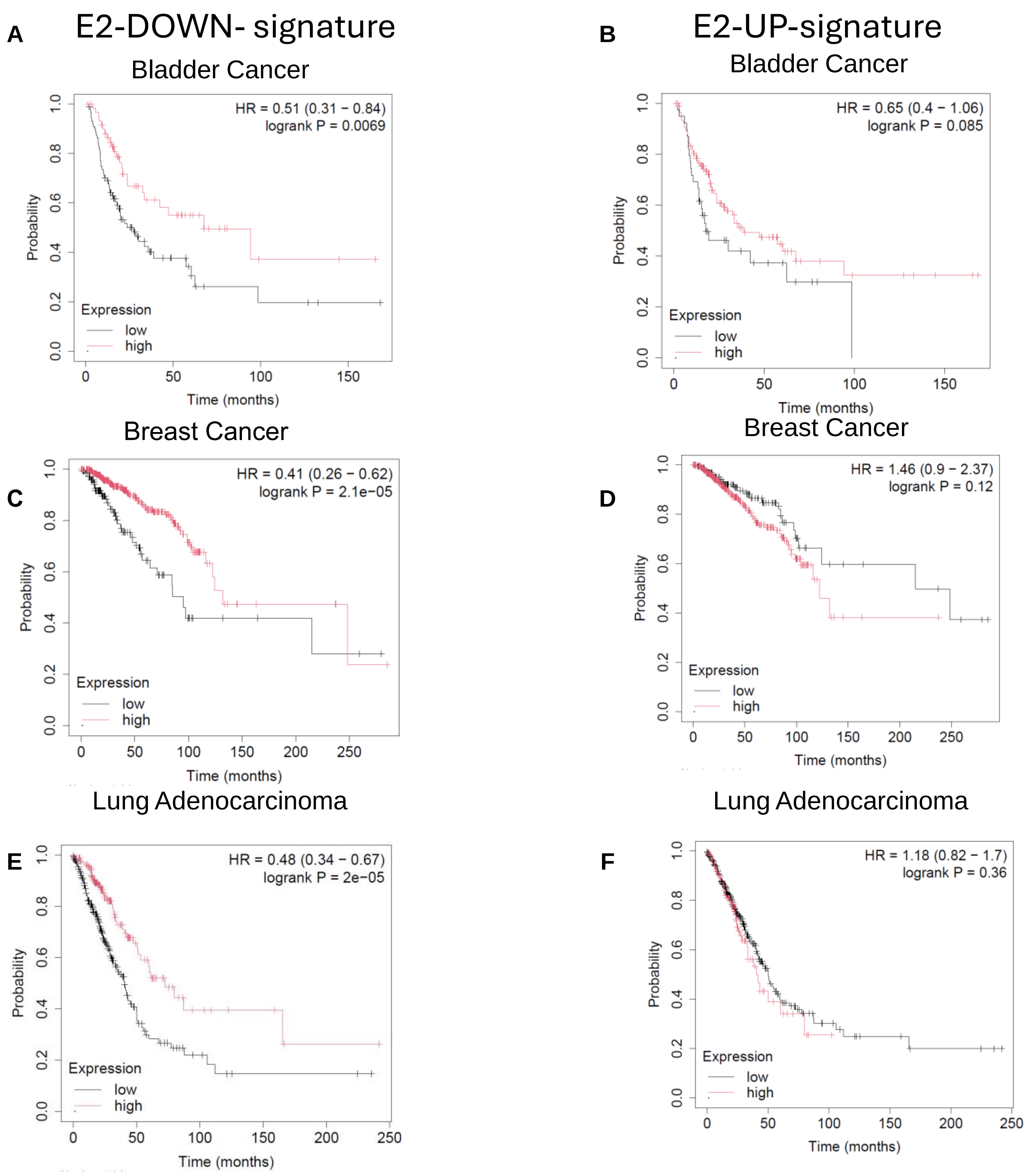

**Supplemental 2 continued: Myeloid Cell Intrinsic Esr1 gene signature predicts survival in patients with varied solid tumor types**

E2- DOWN

E2- Up

**G**

Ovarian Cancer

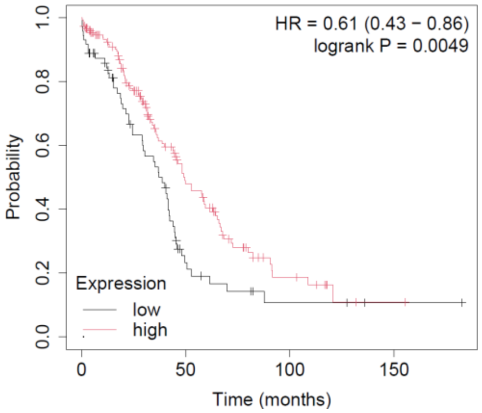


**H**

Ovarian Cancer

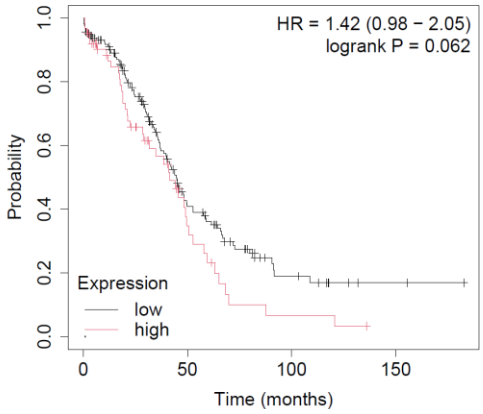


**I**

Sarcoma

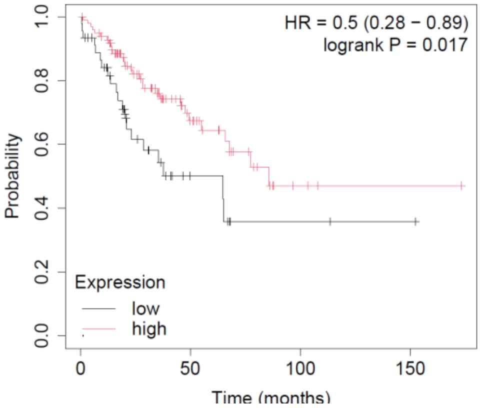


**J**

Sarcoma

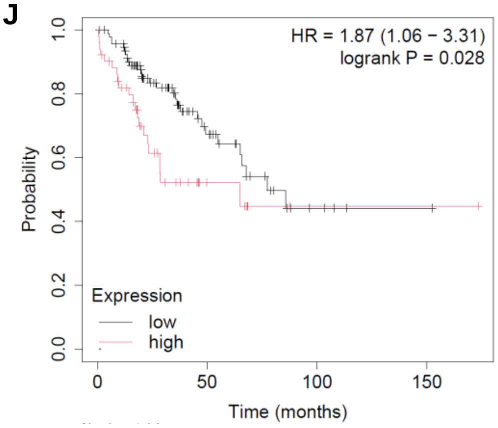


**K**

Uterine corpus endometrial carcinoma

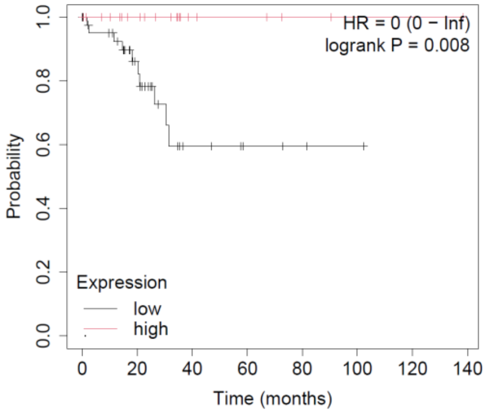


**L**

Uterine corpus endometrial carcinoma

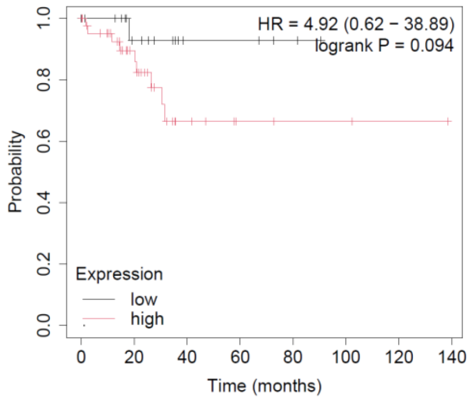

**Supplemental Figure 2. A myeloid cell intrinsic E2 regulated gene signature predicts survival in multiple solid tumor types. Related to Figure 1**: Overall survival in TCGA- bladder cancer (BCa) (**A-B**), breast cancer (BRCA) (**C-D**), lung adenocarcinoma (LUAD) (**E-F**), ovarian cancer (OVC) (**G-H**), sarcoma (**I-J**) and uterine–corpus endometrial carcinoma (**K-L**) using KM Plotter (KM plotter.com)  using mean gene expression signature of bottom 50 ($p<0.05$) or top 50 ($p<0.05$) genes that were significantly changed in intratumoral myeloid cells upon E2 exposure from RNA sequencing.

### Supplemental Figure. 3. Related to Figure 2: E2 suppresses macrophage intrinsic type I IFN signaling only in the presence of apoptotic cells

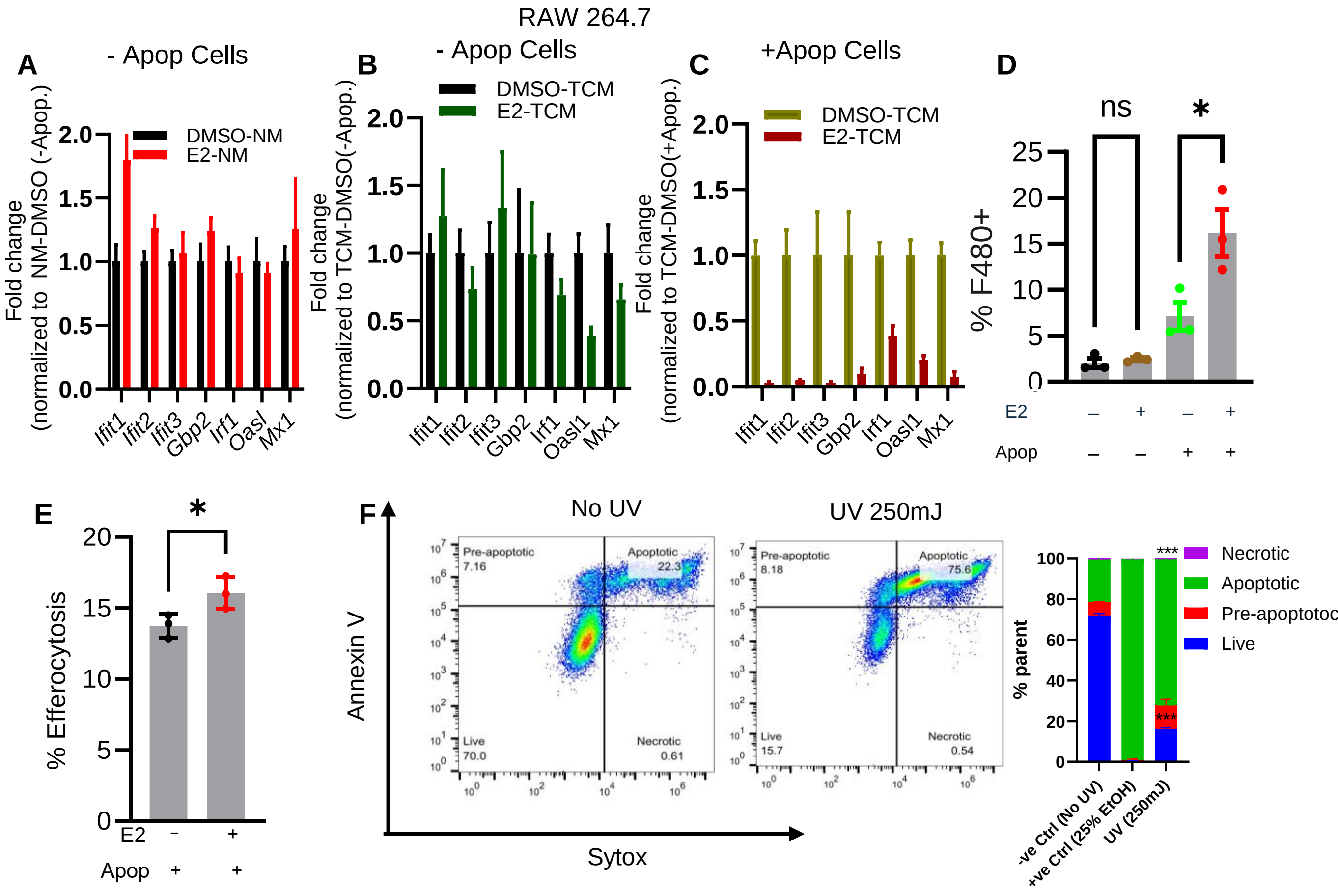


**Supplemental Figure 3. E2 suppresses macrophage intrinsic type I IFN signaling only in the presence of apoptotic cells. Related to Figure 2.** Quantitative PCR of ISGs in RAW264.7 cells that were treated ± E2 in normal media (**A**) ± E2 in tumor-conditioned media (**B**) ± E2 in the presence of apoptotic cells in tumor-conditioned media (**C**). Percentage of mCherry+RAW264.7cells when the cells were treated ± E2 in the presence or in the absence of apoptotic cells (LLC-mCherry-Spectrin) (**D**). Percentage of CTV+ BMDM when the cells were treated ± E2 in the presence or in the absence of apoptotic cells (Jurkat cells) (E). Flow cytometry plots depicting the percentage of apoptotic (Annexin$^{+}$ Sytox$^{+}$) and necrotic cells (Annexin$^{neg}$ Sytox$^{+}$) in Jurkat cells upon U.V treatment (250mJ UV radiation for ~ 3hrs) (F). Percentage of intratumoral MERTK+ TAMs from BPD6 (melanoma) and A7C11 (breast carcinoma) cells injected in ovariectomized mice treated ± E2 Data represent mean ± S.E.M. Significance is calculated by unpaired Students t-test (E), one-way ANOVA (D and E) or two-way ANOVA and pairwise comparison followed by Sidak's multiple corrections (. *p<0.05, **p<0.01, ***p<0.001, ****p<0.0001.)

### Supplemental 4: E2 enhances the expression of CX3CR1 on TAMs

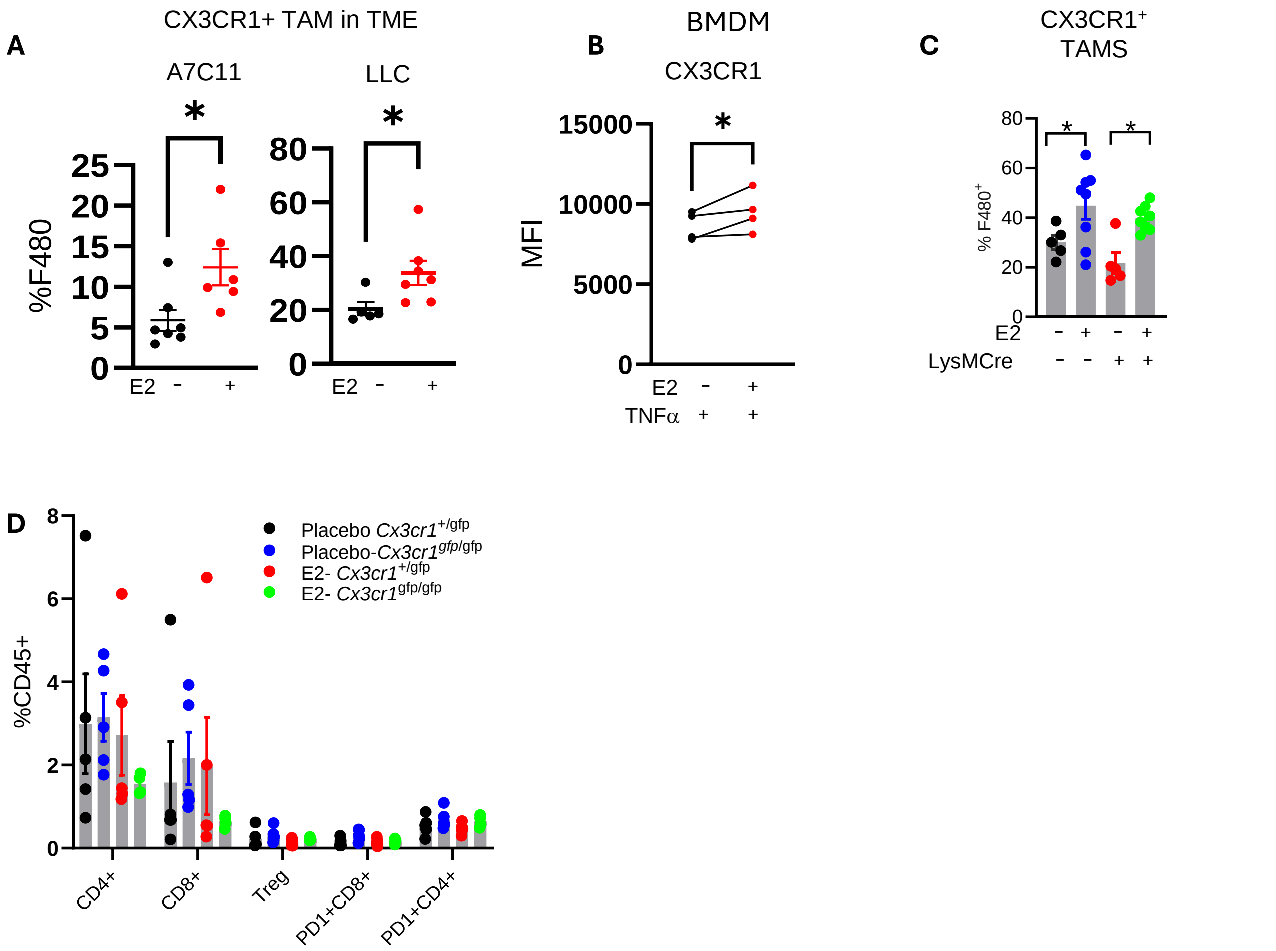


**Supplemental Figure 4**. **E2 enhances the expression of CX3CR1 on macrophages. Related to Figure 4.** Quantification of intratumoral CX3CR1+ TAMs from LLC and A7C11 tumors injected in ovariectomized mice treated ± E2 (**A**). Expression of CX3CR1 in BMDM treated with TNFα (72 hrs) ± E2 (48hrs) (**B**). Quantification of intratumoral accumulation of CX3CR1$^{+}$ TAMs from the experiment described in Figure 3B (**C**). Quantification of intratumoral accumulation of CD4$^{+}$, CD8$^{+}$, Tregs, and exhausted (PD1+) CD4 and CD8+ T cells **(D).** Data represent mean ± S.E.M. Significance is calculated by unpaired Students t-test (A), paired Student's t test (B), one-way ANOVA, and pairwise comparison followed by Sidak's multiple correction (C). #p<0.05 (between CX3CR1 +/gfp - E2 vs placebo, *p<0.05 (CX3CR1gfp/gfp; placebo vs E2) *p<0.05, **p<0.01, ***p<0.001, ****p<0.0001.)

**Supplemental Figure 5: E2 increases NFAT activation to upregulate CX3CR1 expression in tumor-associated myeloid cells.**

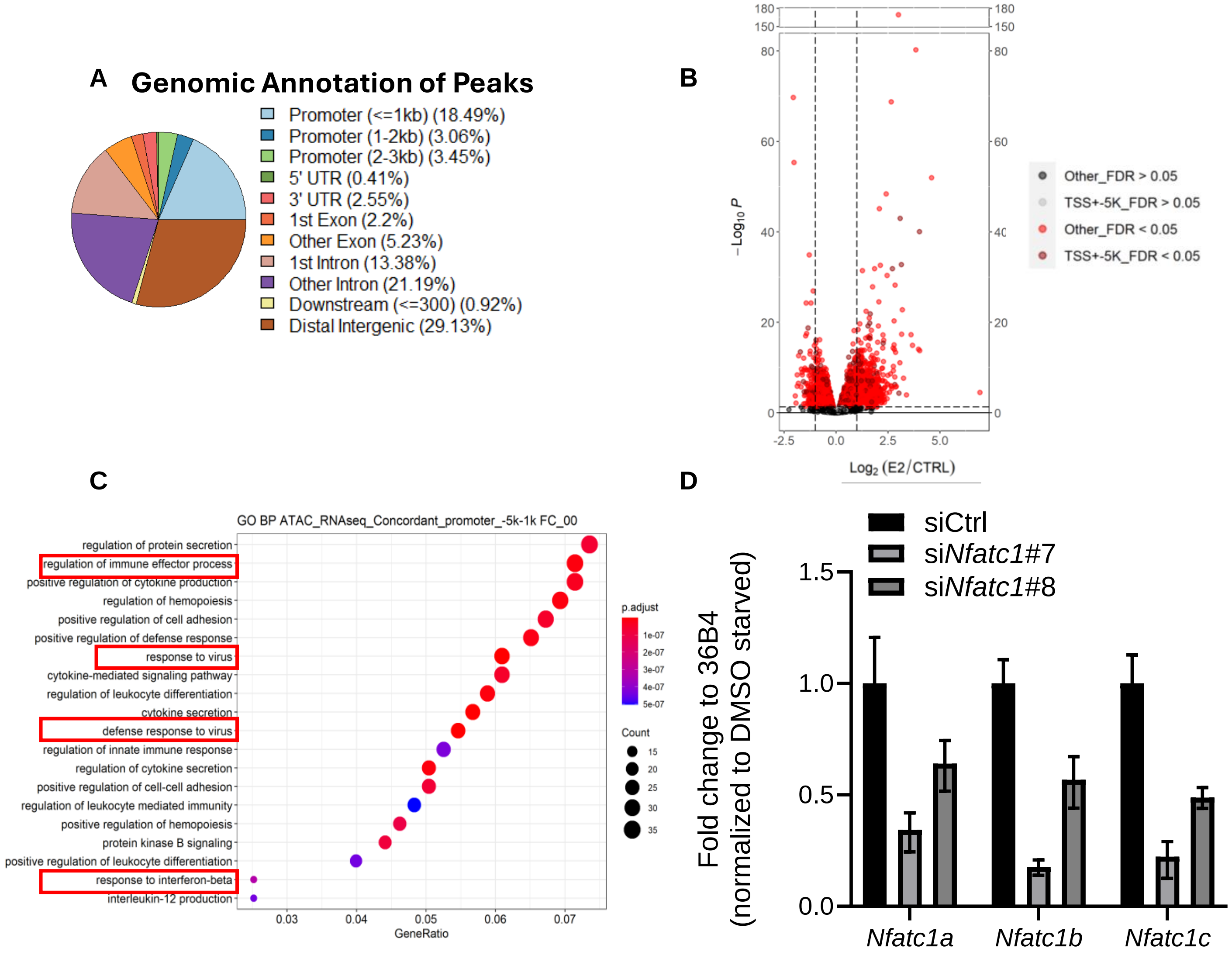


**Supplemental Figure 5**: **E2 increases NFAT activation to upregulate CX3CR1 expression in tumor-associated myeloid cells. Related to Figure 5.** Genomic annotation of significantly enriched differentials peaks upon E2 treatment from ATAC sequencing in tumor-associated myeloid cell from LLC1 tumors treated ± E2 (**A**). Volcano plots showing differentially regulated promoter and promoter-proximal area between placebo and E2 from ATAC-sequencing (**B**). Significantly enriched pathways were computed from combining concordant targets from RNA sequencing and ATAC sequencing (peaks -5k to+1kb promoter sequences) (**C**). Predicted transcription factor binding sites in accessible chromatin from placebo and E2 treated intratumoral (LLC1) myeloid cells; red=predicted TF that acts as a repressor and its expression is decreased (E2 vs placebo) in RNA seq; green=predicted TF that acts as an activator and whose expression is increased in RNA-seq); black=predicted TF that can act as both activator or repressor and whose expression is unchanged (**D**). Gene expression analysis of *Nfatc1a*, *Nfatc1b,* and *Nfatc1c* following siRNA-mediated knockdown of *Nfatc1* in bone-marrow derived macrophages to demonstrate knockdown efficiency.

**Supplemental figure 6: ER modulators enhance efficacy of radiotherapy by suppressing efferocytosis**

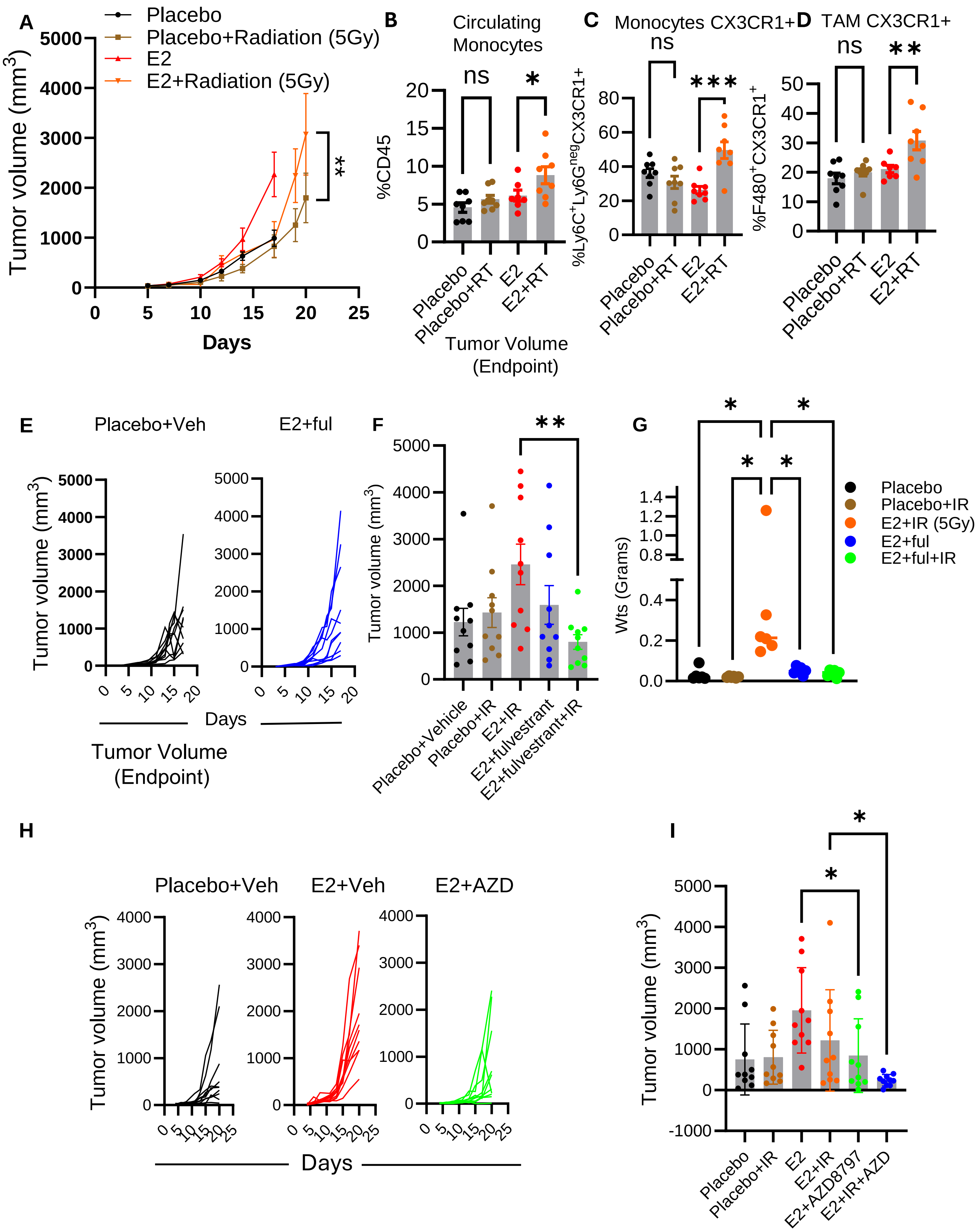

**Supplemental Figure 6. ER modulators enhance efficacy of radiotherapy by suppressing efferocytosis. Related to Figure 7:** Primary tumor growth of LLC cells in ovariectomized mouse treated ± E2 and subjected to radiation (5Gy) (**A**). Flow cytometry analysis and quantification of circulating monocytes, intratumoral CX3CR1+ monocytes and intratumoral CX3CR1 TAMs from LLC tumors treated ± E2 in presence or in absence of ionizing radiation (5Gy) (**B-D**). Tumor growth of LLC cells injected in ovariectomized mice (figure same as 6B) and E2+ful treated group from experiment 6B. (**E**). Final day tumor volume from the experiment described in 6B (**F**). Uterine wet weights from animals from experiment 6B show E2 and fulvestrant exposure (**G**). Tumor growth in control groups from figure 6J (Ovariectomized mice treated with placebo (figure same as 6J), E2 and E2+AZD) (**H**). Final day tumor volume from the experiment is shown in figure 6J (**I**). Significance was calculated by two-way ANOVA and pairwise comparison followed by Dunnett's multiple corrections (), one-way ANOVA and pairwise comparison followed by Sidak's multiple correction (B, C, D and I), pairwise comparison followed by Dunnett's multiple comparisons (F and G). *p<0.05, **p<0.01, ***p<0.001, ****p<0.0001.)

**Supplementary Table I**

Sequences of siRNA

| siRNA | Sequence |
|---|---|
| siNfatc1#7 | UAUACCAGCUCUGCCAUUG |
| siNfatc1#8 | ACGGUUACUUGGAGAAUGA |

**Supplementary Table II.**

Details of qPCR primers (mouse)

| Gene I.D | Forward | Reverse |
|---|---|---|
| *Mx1* | 5’-TGGACATTGCTACCACAGAGGC-3’ | 3’-TTGCCTTCAGCACCTCTGTCCA-5’ |
| *Ifit1* | 5’-CCAAGTGTTCCAATGCTCCT-3’ | 3’-GGATGGAATTGCCTGCTAGA-5’ |
| *Ifit2* | 5’-CTTGACTGTGAGGAGGGGTG-3’ | 3’-TAGTTCGCAATGGCCCATCC-5’ |
| *Ifit3* | 5’-AGACAGGGTGTGCAACCAGG-3’ | 3’-CGACGAATTTCTGATTGATC-5’ |
| *H2q10* | 5’-ACCAGAGTGCAAGTGAGGAGCT-3’ | 3’-GTACTAGGGCAGTGATGTCCTG-5’ |
| *Irf7* | 5’-CCTCTGCTTTCTAGTGATGCCG-3’ | 3’-CGTAAACACGGTCTTGCTCCTG-5’ |
| *Isg20* | 5’-CAATGCCCTGAAGGAGGATA-3’ | 3’-TGTAGCAGGCGCTTACACAG-5’ |
| *Nlrc5* | 5’-CTTCCCGCCTCTCCTTCCACAAT-3’ | 3’-CTCCACCTGCCCACATCCTACCA-5’ |
| *Irf1* | 5’-TCCAAGTCCAGCCGAGACACTA-3’ | 3’-ACTGCTGTGGTCATCAGGTAGG-5’ |
| *Oas1* | 5’-GAGGTGGAGTTTGATGTGCTGC-3’ | 3’-GTGAAGCAGGTAGAGAACTCGC-5’ |
| *Gbp2* | 5’-CTGCACTATGTGACGGAGCTA-3’ | 3’-GAGTCCACACAAAGGTTGGAAA-5’ |
| *Trpv4* | 5’-ACAAGAAGCGCCTGACTGAT-3’ | 3’-GATGAATTCACGCATGTTGC-5’ |
| *Xcr1* | 5’-CCTACGTGAAACTCTAGCACTGG-3’ | 3’-AAGGCTGTAGAGGACTCCATCTG-5’ |
| *Slc25a3* | 5’-CAGCCTGGTTATGCCAACACCT-3’ | 3’-CTGTCTCATCCACAGAGGAGCA-5’ |
| *Cx3cr1* | 5’-CTGTTATTTGGGCGACATTG-3’ | 3’-AACAGATTTCCCACCAGACC-5’ |
| *Rplpo* | 5’-AGATTCGGGATATGCTGTTGGC-3’ | 3’-TCGGGTCCTAGACCAGTGTTC-5’ |

**Supplementary Table III.**

Antibodies used for flow cytometry staining.

| Reagents | Clone | Source | Catalog number | Working concentration |
|---|---|---|---|---|
| Live/Dead Staining dye | | Invitrogen | L34964 | 1:200 |
| BV650 anti mouse CD45 | 30-F11 | Biolegend | 103139 | 1:800 |
| PerCpCy5.5 anti mouse CD3 | 17A2 | BD | 560572 | 1:50 |
| AF647 anti human/mouse Granzyme B | GB11 | Biolegend | 515406 | 1:50 |
| AF700 anti mouse/human CD44 | IM7 | Biolegend | 103026 | 1:100 |
| APC-Cy7 anti hamster CD69 | H1.2F3 | Biolegend | 104526 | 1:100 |
| BV650 anti rat CD8 | 53-6.7 | Biolegend | 100742 | 1:100 |
| BV785 anti rat CD4 | RM4-5 | Biolegend | 100552 | 1:100 |
| BV711 anti rat IFNg | XMG1.2 | Biolegend | 505835 | 1:50 |
| PE anti rat FOXP3 | FJK-16s | eBioscience | 12-5573-82 | 1:100 |
| BV510 anti mouse PD1 | 29F1A12 | Biolegend | 135241 | 1:100 |
| APC anti rat CD25 | PC61.5 | eBioscience | 17-0251-82 | 1:100 |
| PE anti rat CD11b | M1/70 | Biolegend | 101208 | 1:50 |
| AF488 anti mouse CD206 | C06C2 | Biolegend | 141710 | 1:100 |
| BUV496 anti rat CD24 | M1/69 | BD | 564664 | 1:100 |
| PerCPCy5.5 anti mouse monoclonal CD64 | X54-5/7.1 | Biolegend | 139308 | 1:100 |
| APC anti rat F4/80 | BM8 | Biolegend | 123116 | 1:100 |
| APC Cy7 anti hamster CD11c | HL3 | BD | 561241 | 1:50 |
| PE-CY7 anti rat MHCII | M5/114-15.2 | eBioscience | 25-5321-82 | 1: 600 |
| BV711 anti rat Ly6 C | HK1.4 | Biolegend | 128037 | 1:100 |
| PerCPCy5.5 anti mouse CX3CR1 | SA011F11 | Biolegend | 149010 | 1:100 |
| APC Cy7 Anti Mouse MERTK | 2B10C42 | Biolegend | 151520 | 1:100 |
| BV786 anti mouse Ly6 G | 1A8 | Biolegend | 127645 | 1:100 |

**Supplementary Table IV**

Sequence of Barcodes used in ATAC-sequencing

| Sample | Barcode #1 | Barcode #2 |
|---|---|---|
| Placebo-1 | TAGATCGC | GTAGAGGA |
| Placebo-2 | CTCTCTAT | AAGAGGCA |
| Placebo-3 | TATCCTCT | CGAGGCTG |
| Placebo-4 | AGAGTAGA | GCTACGCT |
| E2-1 | GTAAGGAG | CAGAGAGG |
| E2-2 | ACTGCATA | CTCTCTAC |
| E2-3 | AAGGAGTA | TAGGCATG |
| E2-4 | CTAAGCCT | GGACTCCT |
